# Seizures, increased interhemispheric synchrony, altered brain transcriptomics and a leaky blood-brain barrier result from loss of *ap3b2* in a CRISPR tadpole model of DEE48

**DOI:** 10.64898/2025.12.29.696940

**Authors:** Sulagna Banerjee, Cabriana W. Earl, Samuel C. Robson, Paul Szyszka, Caroline W. Beck

**Affiliations:** Department of Zoology, University of Otago, Dunedin, New Zealand; Institute of Life Sciences and Healthcare, School of Medicine, Pharmacy and Biomedical Sciences, University of Portsmouth, PO1 2DT, UK; Centre for Enzyme Innovation, School of Earth and the Environment, University of Portsmouth, PO1 2DT, UK; Genetics Otago Research Centre, University of Otago, Dunedin, New Zealand

**Keywords:** Developmental and Epileptic Encephalopathy, *Xenopus laevis*, GCaMP6s, *ap3b2*, CRISPR, seizure, model organism, GABA pathway

## Abstract

Loss-of-function variants in AP3B2, a neuronal adaptor protein required for synaptic vesicle formation, cause a severe early-onset neurodevelopmental epilepsy known as Developmental and Epileptic Encephalopathy 48 (DEE48). To investigate how AP3B2 loss alters brain development, leading to increased seizure susceptibility, we generated a *Xenopus laevis* model by targeting the orthologous gene using CRISPR/Cas9. *Ap3b2.S*^-^/^-^ (mosaic) F_0_ tadpoles displayed increased locomotor activity with frequent seizure-like episodes when compared to sibling controls. Visualisation of forebrain and midbrain activity using the genetically encoded Ca^2+^ sensor GCaMP6s detected spontaneous, large amplitude, prolonged and widespread neural activity, alongside increased interhemispheric synchrony of both regions. Comparison of whole-brain transcriptomes from *ap3b2* CRISPants and unedited sibling controls detected mainly downregulation of brain expressed genes, with significant over-representation of pathways involved in ion transport, axon formation and guidance, inhibitory (GABA) neurotransmission, and transport across the blood-brain barrier (BBB). In a simple assay for BBB integrity, CRISPant tadpoles were confirmed to have faster leakage of sodium fluoresceinAcute exposure to the angiotensin receptor blocker losartan significantly reduced locomotor hyperactivity, and CRISPant cohorts treated with losartan tended to have lower neural activity, indicating incomplete rescue of the *ap3b2.S* CRISPant phenotype. These findings demonstrate how AP3B2 loss of function alters brain development and the establishment of the BBB, with the resulting alterations in neurotransmitter pathways predisposing the brain to spontaneous seizures. Our results suggest that traditional anti-seizure medications designed to alter ion transport and GABA metabolism could be augmented with drugs targeting neuroinflammation, as adjunct seizure control options in infants with DEE48.

## 1 Introduction

Developmental and epileptic encephalopathies (DEEs) are a heterogeneous group of severe, early-onset epilepsies that arise from genetic abnormalities and profoundly affect neurodevelopment. They are characterized by frequent, often refractory seizures, accompanied by global developmental delay, cognitive impairment, and behavioural regression (Scheffer and Liao, 2020, Scheffer et al., 2024). A defining feature is the association between epileptiform EEG activity and developmental regression, underscoring the dynamic interplay between seizure activity and brain maturation (Scheffer et al., 2024). Advances in molecular genetics have revealed that DEEs reflect perturbations across diverse neuronal and developmental pathways. According to the Online Mendelian Inheritance in Man catalogue (OMIM, https://www.omim.org), 119 genes are currently recognized as causative, but emerging studies suggest that variants in more than 800 genes may contribute to the broader DEE spectrum (Poke et al., 2023). These discoveries link DEEs to both neurodevelopmental and neurodegenerative mechanisms, offering new insight into disease pathogenesis (Riva et al., 2025). Although individual DEE syndromes are rare, their collective incidence approaches 1 in 590 children, making them a major cause of early-life neurological disability (Poke et al., 2023, Symonds et al., 2021).

Among the expanding number of genes implicated in DEEs, *AP3B2* exemplifies how disruption of synaptic vesicle trafficking can result in severe and early neurodevelopmental failure. Biallelic pathogenic variants in *AP3B2* were first identified by Assoum et al. through whole-exome sequencing of individuals with early-onset epileptic encephalopathy, defining developmental and epileptic encephalopathy-48 (DEE48 OMIM#617276) (Assoum et al., 2016). Analysis of a cohort of 12 individuals with DEE48 from 8 families revealed three nonsense mutations (p.Arg67*, p.Glu152* and p.958*), two frameshift mutations, each -4 bp (p.Leu841Glnfs*10 and p.Thr1060Ser*7), and three splice variants deleting exons 10 and 14 (Assoum et al., 2016) (Figure1). Seizure onset occurred within the first year of life, ranging from birth to early infancy, and included infantile spasms, myoclonic seizures, and generalized or focal seizure types, frequently accompanied by markedly abnormal EEG patterns such as hypsarrhythmia. Neurodevelopmental impairment was profound and often evident prior to or independent of seizure onset, with severe hypotonia, absent or minimal speech, poor visual engagement frequently associated with optic atrophy, and postnatal microcephaly, while early brain MRI findings were often unremarkable (Assoum et al., 2016).

Subsequent case series have reinforced the consistency and severity of the DEE48 phenotype. Two further individuals with homozygous truncating frameshift variants in *AP3B2* (p.Ala149Serfs*34 and p.Pro993Argfs*5) presented with refractory seizures beginning at approximately 3–4 months of age, severe global developmental delay, hypotonia, stereotypic movements, postnatal microcephaly, and progressive intellectual disability (Dilber et al., 2022). Genomic analysis of a cohorts enriched for early-onset epilepsy and intellectual disability, identified two cases of the frameshift p.Glu613Ser*182 (Anazi et al., 2017). More recently, a novel homozygous *AP3B2* missense variant (p.Val106Ile) was identified in a child with neonatal-onset seizures, microcephaly, developmental regression, and electroclinical features consistent with DEE48, extending the allelic spectrum while preserving the core clinical phenotype (Alizadeh et al., 2025). Together, these studies show that homozygous loss of function of AP3B2 causes DEE.

AP3B2 encodes the β3B subunit of the neuronal adaptor-protein-3 (AP-3) complex, which mediates cargo selection and vesicle budding from endosomes (Dell’Angelica et al., 1997, Faúndez et al., 1998). The neuronal AP-3 complex, enriched in axons and presynaptic terminals, is essential for synaptic vesicle biogenesis (Blumstein et al., 2001). Loss of AP3B2 disrupts the targeting of key vesicular proteins such as ZnT3 and ClC-3, causing altered vesicle composition and reduced vesicular zinc content, which compromises synaptic transmission (Seong et al., 2005) and leads to hyperactivity, seizures, and synaptic dysfunction in *Ap3b2*⁻/⁻ mice (Nakatsu et al., 2004). Subsequent studies have expanded the *AP3B2* mutational spectrum, identifying novel homozygous and compound heterozygous loss-of-function (LOF) variants (Figure 1) associated with autosomal-recessive DEE48 (Anazi et al., 2017, Dilber et al., 2022, Alizadeh et al., 2025).

**Figure 1.**
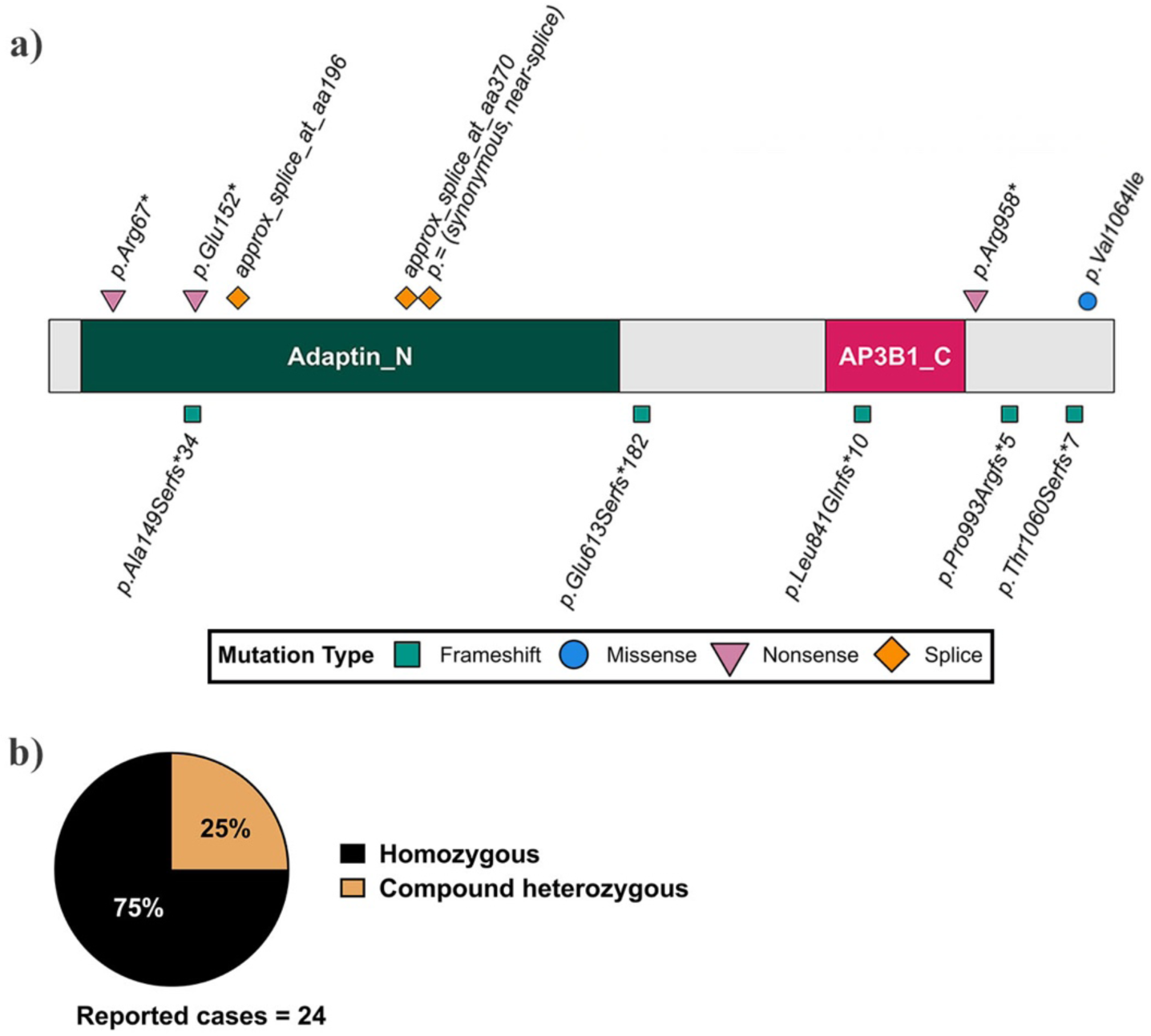
Summary of reported AP3B2 mutations and inheritance patterns in patients. (a) Schematic representation of the human AP3B2 protein showing the locations and types of reported pathogenic variants. The conserved Adaptin N terminal (Adaptin_N) and Clathrin-adaptor complex-3 beta-1 subunit C-terminal (AP3B1_C) domains are indicated, with frameshift (green squares), missense (blue circle), nonsense (pink triangles), and splice-site (orange diamonds) mutations annotated along the protein. (b) Summary of reported inheritance patterns across 24 documented cases, 75% were homozygous for AP3B2 variants, while 25% are compound heterozygous. The DEE48 pathogenic variants in *AP3B2* shown here were described in Assoum et al. (2016), Anazi et al. (2016), Dilber et al. (2022) and Alizadeh et al. (2025).

Despite rapid progress in genetic diagnosis, effective therapies for DEEs remain limited, highlighting the need for experimental models that permit direct observation and manipulation of early neurodevelopmental processes in vivo. To address this, simple vertebrate systems such as the *Xenopus* tadpole have emerged as powerful platforms for functional analysis of DEE-associated genes. Early chemoconvulsant-based *Xenopus laevis* seizure models enabled electrophysiological and behavioural quantification of seizure activity (Hewapathirane et al., 2008, Panthi et al., 2024, Bell et al., 2011). More recently, CRISPR/Cas9-mediated gene editing has allowed creation of tadpole models of NeuroD2 haploinsufficiency in both *X. laevis* and *X. tropicalis* that recapitulate the phenotype of DEE72, facilitating rapid assessment of pathogenic mechanisms and therapeutic responses (Banerjee et al., 2024, Sega et al., 2019).

To investigate the aetiology of DEE48 and further understanding of how loss of function mutations in AP3B2 increase seizure susceptibility, we generated a *Xenopus laevis* tadpole model by targeting the orthologous gene using CRISPR/Cas9. *Xenopus* are well suited for this purpose, due to efficient CRISPR introduction of disruptive frameshift indels, easy access to the developing tadpole brain and a drug permeable skin (Li et al., 2022, Banerjee et al., 2024).We mined data from a previous study of brain transcriptomics in this species (Ta et al., 2021) and found that *ap3b2.S* is expressed in the developing tadpole brain. *Ap3b2.S*^-^/^-^ (mosaic) CRISPants replicated the seizure phenotype previously shown in the mouse model (Nakatsu et al., 2004). These tadpoles showed seizure-like behaviour, defined as increased mean swimming velocity and runs of C-starts with abrupt directional changes (darting), compared to unedited sibling controls.GCaMP6s imaging of *ap3b2* CRISPant tadpoles revealed increased frequency and amplitude of Ca²⁺ events with increased cross-regional synchrony, consistent with hypersynchronous epileptic activity. Despite a grossly normal brain morphology, sodium fluorescein tracking demonstrated early and pronounced blood-brain barrier (BBB) leakage. Transcriptomic analysis of the tadpole brain showed prominent dysregulation of both neuronal signalling and development pathways as well as evidence of altered neuroinflammatory markers. Losartan, an angiotensin-II receptor blocker previously shown to reduce seizures in NeuroD2 (DEE72) CRISPants (Banerjee et al., 2024), significantly suppressed swimming velocity. Collectively, these findings establish the DEE48 *X. laevis* CRISPant tadpole as a rapid, scalable, and physiologically relevant vertebrate model for dissecting *AP3B2*-associated epileptic encephalopathy and evaluating targeted interventions.

## 2 Materials and methods

### 2.1 Production and maintenance of *Xenopus laevis* embryos

Adult *Xenopus laevis* frogs were maintained in temperature-controlled aquaria under standard husbandry conditions and handled in accordance with institutional animal ethics requirements. All procedures were approved by the University of Otago Animal Ethics Committee under protocols AUP22/12 and 22/24. To induce ovulation, adult females were primed by injection of human chorionic gonadotropin (hCG; 500 IU per 75 g body weight) into the dorsal lymph sac 16 hours before egg collection. Primed females were housed overnight in pairs in small holding tanks containing “frog water” (carbon-filtered tap water). Once egg laying commenced, each female was transferred to individual tanks containing 1 L of 1× Marc’s Modified Ringers (MMR; 10 mM NaCl, 0.2 mM KCl, 0.1 mM MgSO₄·6H₂O, 0.2 mM CaCl₂, 0.5 mM HEPES, 10 µM EDTA, pH 7.8), and eggs were collected hourly. Eggs were fertilized in vitro using a sperm suspension prepared from freshly isolated testes of a euthanized adult *Xenopus laevis* male. Fertilized eggs were left undisturbed until embryo rotation occurred (15–20 min), then dejellied in 2% L-cysteine (pH 7.9). Embryos were maintained at 14–18 °C, monitored regularly, and staged according to the Nieuwkoop and Faber (NF) normal table (Nieuwkoop and Faber, 1994).

### 2.2 CRISPR/Cas9-mediated targeting of *ap3b2.S*

Using published transcriptomic datasets (Ta et al., 2021), the *ap3b2.S* homeologue was confirmed to be expressed in *X. laevis* brain tissue and identified as the ortholog of human AP3B2. Guide RNAs targeting exonic regions of *ap3b2.S* were designed using ChopChop (Labun et al., 2016) (https://chopchop.cbu.uib.no/) and evaluated in InDelphi (Shen et al., 2018) (https://indelphi.giffordlab.mit.edu/) to confirm predicted on-target efficiency and minimize off-target editing (Supplementary figures S1–S2). Two sgRNAs, ChopChop rank 105 (sgRNA2) and rank 84 (sgRNA3), were selected to disrupt *ap3b2.S* (henceforth described as *ap3b2*). The sgRNA sequences and PCR primer sets used for genotyping are listed in Table 1. A scrambled control sgRNA (CTTGTAGATCAGGTGCAAGCTGG) was designed using the method of (Hsu et al., 2013). BLAST analysis confirmed that this scrambled sequence had no predicted targets in the *X. laevis* genome. For sgRNA synthesis, long 54–55 bp oligonucleotides were designed using the EnGen sgRNA Template Oligo Designer (NEB), incorporating the 20-nt ChopChop target sequence (excluding the NGG PAM). A 5′ G was added when absent to optimize transcription efficiency. Oligonucleotides were synthesized by IDT and used with the EnGen sgRNA Synthesis Kit, S. pyogenes (NEB). Synthesized sgRNAs were dissolved in nuclease-free water, aliquoted, stored at -80 °C, and thawed on ice prior before injections. Immediately prior to use, 0.3 µL of EnGen *S. pyogenes* Cas9-NLS protein was added to the sgRNA and incubated at 37 °C for 5 minutes to form Cas9–sgRNA ribonucleoprotein complexes.

**Table 1.**
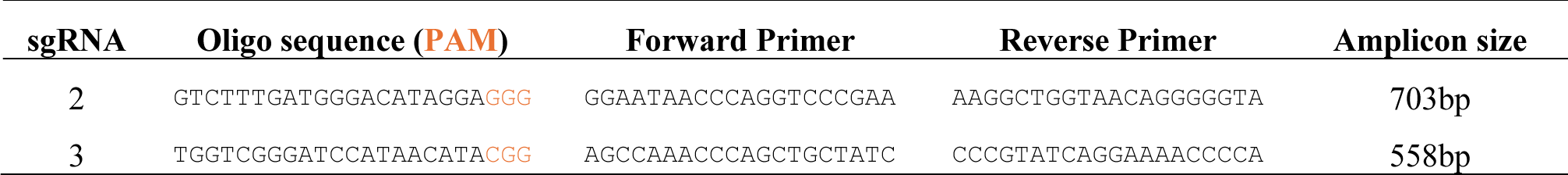
*ap3b2.S* sgRNAs and primer sequences.

Fertilized, de-jellied embryos (<1 hour post-fertilization) were examined for sperm entry points and aligned in rows within a 2 mm × 40 mm trench cut into a 50 mm agar-coated dish filled with 5% Ficoll PM400 in MMR. Cas9–sgRNA complexes were back-filled into pulled Drummond glass needles and bilateral 5 nL injections (total 10 nL per embryo) were delivered adjacent tothe female pronucleus using a Nanoject II injector mounted on a Narishige MM-3 micromanipulator. sgRNA doses were 300 -350 pg per embryo. Embryos were injected in batches of 25 and transferred immediately to 24 °C. For each sgRNA, 100 embryos from the same sibship were injected. Control embryos were injected with Cas9-NLS protein pre-incubated with the scrambled sgRNA in the same amounts. At 2-3 hours post-injection, embryos were assessed for normal cleavage and transferred to 2.5% Ficoll in 0.1× MMR. The following day, embryos were re-screened for normal development, and five were randomly selected for genotyping. Remaining embryos were maintained in 0.1× MMR until stages 46–47. For genotyping, whole embryos or individual tadpoles were homogenized in 200 µL of 5% Chelex in PBS with 1.5 µL Proteinase K (25 mg/mL) and incubated at 65 °C for 3 hours (embryos) or overnight (tadpoles). Reactions were terminated at 95 °C for 5-10 minutes and briefly centrifuged to pellet the Chelex resin. One microlitre of supernatant was used directly as PCR template (primer sequences in Table 1). Gene editing was confirmed using TIDE analysis (Brinkman et al., 2014) (https://tide.nki.nl/), comparing CRISPant samples to scrambled sgRNA controls, and validated against InDelphi predictions.

### 2.3 Behavioural recording and TopScan-based locomotor analysis

Behavioural recordings were performed using a high-resolution locomotor tracking protocol developed for *Xenopus laevis* tadpoles, adapted from the automated TopScan analysis workflow described in Banerjee et al. (2024). Stage 47 tadpoles were placed individually into wells of a clear 24-well plate containing 0.1× MMR and allowed to acclimate for 2–3 minutes before recording. Plates were positioned on a uniform full spectrum daylight LED back-illumination stage, and overhead recordings were acquired using two identical 12.3 Megapixel camera with 16 mm telephoto lens mounted at a fixed height and controlled by a Raspberry Pi5.. Tadpoles in each 24-well plate was recorded for a continuous 1 hour period at 50 frames per second under constant illumination. Raw video files were converted to .mp4 format using HandBrake (https://handbrake.fr/) to standardize encoding prior to analysis. Automated locomotor quantification was conducted in TopScan (CleverSys Inc.) following the arena-based workflow optimized for *X. laevis* tadpoles (Panthi et al., 2024). For each video, a static background image was generated, and circular arenas corresponding to each well were manually defined. Identical detector thresholds, background parameters, and arena definitions were applied across all recordings to ensure analytical consistency.

Locomotor metrics were extracted using the TopScan LocoMotion Super module, including total distance travelled and mean swim velocity (mm/sec). Darting behaviour was quantified using the DrugAbuse detector, with darting defined as rapid burst movements exceeding the pre-set high-velocity threshold. Event durations, velocities, and locomotor trajectories were exported as summary tables for downstream statistical analysis. All recordings were processed using identical acquisition and analysis settings to ensure reproducibility across biological replicates.

### 2.4. *In vivo* Ca²⁺ imaging and analysis

*In vivo* Ca²⁺ imaging was performed using a widefield fluorescence protocol adapted from the *neuroD2* DEE72 *Xenopus laevis* tadpole model and a previously published cranial-window workflow (Banerjee et al., 2024). Single-cell embryos were bilaterally injected with 500 pg GCaMP6s and 250 pg mCherry mRNA together with the respective Cas9/sgRNA reagents. At NF stage 47, tadpoles were anaesthetized in 1:4,000 MS-222 for 2 minutes, positioned dorsal-side-up in a Petri dish, and embedded in 2% low-melting-point agarose. Embedded animals were submerged in 0.1× MMR containing 200 µM pancuronium bromide to ensure complete neuromuscular blockade. A cranial window was created by gently removing the dorsal head skin with fine forceps to expose the dorsal brain surface, which was stabilised under a thin cap of 1% low-melting-point agarose (Supplementary figure S3). Tadpoles were imaged using a Zeiss Axio Examiner D1 upright fluorescence microscope equipped with a 10×/0.3 NA water-dip objective (N-Achroplan, Zeiss). GCaMP6s fluorescence was excited at 480 nm using a Polychrome V light source (TillPhotonics) and detected through a 495 nm dichroic mirror and a 505 nm long-pass emission filter. Images were acquired with a PCO Sensicam CCD camera using 4×4 on-chip binning, as described in Banerjee et al (2024). Imaging was restricted to the forebrain and midbrain, as inclusion of the hindbrain reduced the effective field of view and compromised spatial resolution and signal quality in these anterior regions. Spontaneous brain activity was recorded for 30 minutes at 2 frames/s (400 ms exposure per frame). Raw image sequences were processed in Fiji (Schindelin et al., 2012) to generate ΔF/F₀ (%) stacks. For each recording, a median Z-projection across the full 30-minute stack was used to compute the baseline image (F₀); ΔF was calculated as F(t) − F₀, and ΔF/F₀ values were expressed as percentages (ΔF/F₀%).

ΔF/F₀% image stacks were analysed in MATLAB. A whole-brain mask was first manually delineated on the maximum-fluorescence projection using interactive freehand drawing tools (imfreehand) and applied across the full image stack to restrict analysis to brain tissue. CalciSeg (Günzel et al., 2024) was then applied within this masked region for automated, correlation-based detection and refinement of Ca^2+^-active regions of interest (ROIs), as well as for initial preprocessing; all subsequent signal extraction and quantitative analyses were performed in MATLAB (MathWorks). For each ROI, a binary mask was applied to every frame of the ΔF/F₀% stack. At each time point, ΔF/F₀% pixel values within the ROI were extracted and averaged to yield a single fluorescence intensity value. Repeating this operation across frames generated ΔF/F₀% traces for each ROI, which was used for event detection and downstream analyses. Initial denoising occurred implicitly during CalciSeg processing, which suppresses pixel-level noise and background fluctuations by retaining spatially and temporally correlated Ca^2+^signals. No additional spatial or temporal denoising filters were applied prior to event detection.

To remove slow baseline drift, ΔF/F₀% traces were high-pass filtered at 0.005 Hz. For event detection, a single fixed threshold was derived from control animals by pooling high-pass–filtered whole-brain mean traces and defining the cutoff as 3× the standard deviation (SD) of this pooled control distribution. This control-derived threshold was applied uniformly to all replicates and experimental groups; thresholds were not calculated separately for individual traces. For each tadpole, fluorescence values were averaged across all pixels within the manually defined whole-brain mask at each time point to generate a single whole-brain mean ΔF/F₀% trace. Peaks were identified on the high-pass–filtered whole-brain trace using minimum width and distance constraints (5 s each), and peaks exceeding the fixed control-derived threshold were classified as Ca^2+^ events.

For frequency-domain analysis, whole-brain ΔF/F₀% traces were subjected to fast Fourier transformation (FFT) on a per-ROI basis. Single-sided FFT amplitudes were squared to obtain power spectral density estimates ((ΔF/F₀)²/Hz), interpolated onto a common 0.01–1 Hz frequency grid, and averaged across Ca²⁺-active ROIs to generate a whole-brain spectrum per tadpole. Group spectra are shown as mean power ±95% confidence intervals across animals. Integrated low-frequency power (0.01–1 Hz) was calculated as the area under the power spectrum.

To quantify neural synchrony between brain regions, Pearson correlation coefficients were calculated between ΔF/F₀% traces of the left and right forebrain and left and right midbrain. To minimise light scatter from neighbouring regions, signals were extracted from fixed-area (500-pixel) central ROIs positioned at the geometric centroids of each region. Forebrain and midbrain boundaries were manually delineated on maximum-fluorescence images split into left and right hemispheric masks, and centroids were computed to define circular ROIs of 500 pixels(regionprops, Centroid).

### 2.5 Losartan treatment for behavioural and Ca^2+^ imaging assays

#### Behavioural Phenotyping Following Losartan Treatment

Losartan treatment assays were conducted using stage 47 *Xenopus laevis* tadpoles generated as described above. Individual tadpoles were placed into wells of a clear 24-well plate containing 0.1× MMR and allowed to acclimate for 2–3 minutes before baseline recording. Baseline locomotor activity for each *ap3b2.S* CRISPant was recorded for 1 hour using the Raspberry Pi high-resolution behavioural tracking setup described in Section 2.3. Following the baseline recording, 200 µL of 50 mM losartan (Sigma) dissolved in MilliQ water (MQW) was added directly to each well, resulting in a final concentration of 10 mM losartan. Tadpoles were incubated in Losartan for 1 hour under identical environmental conditions. Immediately after incubation, a second 1-hour behavioural recording was performed using the same imaging setup and acquisition parameters. Raw video files were converted to .mp4 format and analysed in TopScan (CleverSys Inc.) using the same detector thresholds, arena definitions, and workflow described in Section 2.3. The LocoMotion Super and DrugAbuse (darting) detectors were used to extract locomotor and high-velocity event metrics. Behavioural parameters recorded before and after Losartan treatment were compared within the same CRISPant tadpoles to assess drug-induced changes.

#### Ca^2+^ Imaging Following Losartan Treatment

To assess the effect of losartan on neuronal activity, in vivo Ca²⁺ imaging was performed on tadpoles treated with losartan or left untreated. CRISPants in the control group received no drug exposure and were imaged using the standard cranial-window GCaMP6s protocol described in Section 2.4. For the treated group, *ap3b2* CRISPants were incubated in 10 mM losartan for 1 hour immediately prior to cranial-window preparation. Following incubation, tadpoles were prepared for in vivo Ca²⁺ imaging as above.

### 2.6 Blood–brain barrier permeability assay

BBB permeability was assessed using a modified sodium fluorescein (NaF) leakage assay adapted from the *neurod2* DEE72 *X. laevis* tadpole model (Banerjee et al., 2024). NF stage 47 tadpoles were positioned in a Petri dish and embedded in 2% low-melting-point agarose. Once the agarose had set, agarose-embedded animals were submerged in 1:4,000 MS-222 in 0.1× MMR for the duration of the experiment. NaF dye (10 nL of a 0.1 mg/mL solution) was then injected into the fourth ventricle using a glass capillary needle following the same injection approach used for embryos. Following injection, tadpoles were kept protected from light. Dorsal images of the head were acquired at 2, 5, 10, and 20 minutes post-injection using a Leica Fluo III stereomicroscope with a GFP2 filter set and a DFC7000T camera under fixed exposure settings. Images were analyzed offline in Fiji. Mean fluorescence intensity (MFI) was extracted from the green channel within a defined region of interest (ROI). To ensure that measurements reflected BBB permeability rather than injection artefacts, the ROI was always drawn on the side opposite to the injection site, capturing NaF dispersion outside the brain parenchyma.

### 2.7 Transcriptomic sample preparation, RNA sequencing, and bioinformatic analysis

NF stage 47 *Xenopus laevis* tadpoles were anesthetized in a 1:4,000 MS-222 solution until unresponsive to touch, then positioned dorsal-side-up on a custom agar dissection plate. Using fine forceps for stabilization, the brain-spinal cord junction was severed with Vanna iridectomy scissors, and the entire brain (forebrain, midbrain, and hindbrain) was excised as an intact unit by cutting along the cranial margins. For each biological replicate, six brains were immediately pooled into pre-labelled tubes on dry ice and stored at −80 °C. Total RNA was extracted using the RNeasy Mini Kit (Qiagen) with on-column DNase digestion, and purified RNA was stored at −80 °C until sequencing. RNA quality control and library preparation were performed by the Otago Genomics Facility, where high-quality samples were used to generate TruSeq stranded mRNA libraries. Indexed libraries were then pooled and sequenced (Illumina Nextseq 2000 P3-200 XLEAP kit, 2 x 100bp paired end) to a depth of 50-60 million reads per sample.

Raw reads underwent standard quality control, adapter trimming, and alignment to the *X. laevis* v10.1 reference genome (XENLA_10.1 accessed May 2025, Xenbase.org) using STAR (Dobin et al., 2013) at the University of Portsmouth, UK. Sorted BAM files were used to generate a read count matrix in Galaxy.eu using FeatureCounts (Liao et al., 2014) (stranded_reverse, gene_id, paired count as one). Differentially expressed genes were identified from normalised read counts, filtered to remove low or non-expressed gene ids (CPM <=1 in 4 or more samples) using EdgeR (Robinson et al., 2010). Genes with a false discovery rate (Benjamini-Hochberg corrected FDR) < 0.05 and a fold change of >2 were considered differentially expressed between control and ap3b2 CRISPant brains. Sequencing data and read counts are available at NCBI GEO under accession GSE312492

Principal component analysis (PCA) plot was generated in R v4.5.1 from normalised read counts (Log2 counts per million) using FactoMineR (Lê et al., 2008) and factoextra packages (Kassambara and Mundt, 2020) and plots rendered with ggplotR (Wickham, 2016). Volcano plots were generated in R from edgeR statistics. Functional enrichment of differentially expressed genes was performed using Enrichr (https://maayanlab.cloud/Enrichr/) (Chen et al., 2013, Kuleshov et al., 2016, Xie et al., 2021) against a custom background list of brain-expressed genes (Supplementary File 1), incorporating Gene Ontology (Biological Process and Molecular Function) and KEGG pathway libraries. KEGG pathway visualisation was carried out using Pathview, and enrichment results were interpreted alongside ClinVar 2025 annotations to identify overlap with known DEE- and seizure-associated genes. Heatmaps were made from Z-scores of normalised read counts using the pheatmap package (Kolde, 2025) in R v4.5.1 (RDevelopmentCoreTeam, 2024).

### 2.8 Graphs and statistical analysis

Most graphs and statistical analyses were prepared in GraphPad Prism v10. Raw data, normality test results and analysis for all figures can be found in Supplementary File 1. The Shapiro-Wilk test for normality was used to confirm normal distributions, with nonparametric tests being used if distributions were non-normal. Specific tests are included in figure legends.

## 3 Results

### 3.1 CRISPR/Cas9-mediated editing of *ap3b2.S* induces hyperactivity and seizure-associated behaviour

Chemoconvulsant models have established that PTZ-evoked seizures in *X*. *laevis* tadpoles manifest as rapid, uncontrolled tail bends, excessive turning, and recurrent C-shaped body contractions, providing robust behavioural markers of seizure onset and severity (Bell et al., 2011, Hewapathirane et al., 2008, Panthi et al., 2024). Functional DEE models show similar behavioural patterns, as observed in the *neurod2* CRISPant tadpoles, where spontaneous seizure-like behaviours from early larval stages, including abrupt C-shaped convulsions and sustained periods of high-intensity, uncoordinated swimming were reported in both *X. laevis* and *X. tropicalis* models (Banerjee et al., 2024, Sega et al., 2019). We used the same CRISPR knockdown strategy to determine whether disruption of *ap3b2.S* in *X. laevis* reproduces the phenotype of DEE48.

Since DEE48 results from homozygous loss of AP3B2, two sgRNAs were designed and tested to determine which would produce the greatest disruption of *ap3b2.S*. Preliminary behavioural screening revealed that *ap3b2* CRISPants generated with sgRNA2 exhibited markedly elevated swim velocity, increased darting behaviour, and prolonged circling compared with both GFP and Cas9-only controls, whereas sgRNA3 CRISPants showed a more variable phenotype (Supplementary figure S4). Consistent with these behavioural differences, sgRNA2 also produced higher mean editing (85.6% +/-3.5) (Supplementary figure S4). For these reasons, sgRNA2 was selected for all subsequent analyses.

To investigate whether loss of AP3B2 function contributes to DEE-associated behavioural abnormalities, we generated *ap3b2* CRISPants using sgRNA2. The sgRNA2 cut site lies immediately upstream of the AP3B1_C domain, and Sanger sequencing of CRISPant tadpoles confirmed substantial sequence degradation at the targeted locus (Figure 2a). TIDE analysis revealed a heterogeneous mixture of frameshift and in-frame alleles across *ap3b2* CRISPants, with three predominant deletion events: a 7 base pair (bp) frameshift deletion detected in 67% of sequenced samples, an 11 bp frameshift deletion in 83%, and a 12 bp in-frame deletion in 93% of samples (Supplementary figure S5). The 7 bp and 11 bp deletions are predicted to introduce premature stop codons shortly downstream of the cut site, resulting in truncated proteins lacking the conserved AP3B1_C domain (Figure 2b). In contrast, the 12 bp deletion removes four amino acids and would result in an almost full length protein. However, this in-frame deletion lies within the conserved AP3B1_C domain. Since DEE48 patients have seizures from early infancy, we next examined whether disruption of Ap3b2 in tadpoles leads to abnormal locomotor activity, indicative of spontaneous seizures. In the *neurod2* DEE72 tadpole model, seizure-associated neural activity was captured by two characteristic behavioural signatures: darting, defined as rapid, high-velocity C-shaped contractions occurring in abrupt bursts, and elevated mean swimming velocity (Banerjee et al., 2024). High-speed video recordings captured clear darting episodes in *ap3b2* CRISPants (Figure 2c,; <u>Supplementary video 1)</u>. Quantitative tracking using TopScan confirmed that the *ap3b2* CRISPant group had significantly higher mean swimming velocities (Figure 2d) and increased time spent darting compared with sibling controls (Figure 2e). Mean editing in 7 randomly picked embryos from the cohort was 83.3% +/- 3.9 (Figure 2f). These levels of *ap3b2* editing, while not a complete knockout, are therefore sufficient to elicit spontaneous seizure-like locomotor behaviour in mosaic F_0_ *X. laevis* tadpoles.

**Figure 2.**
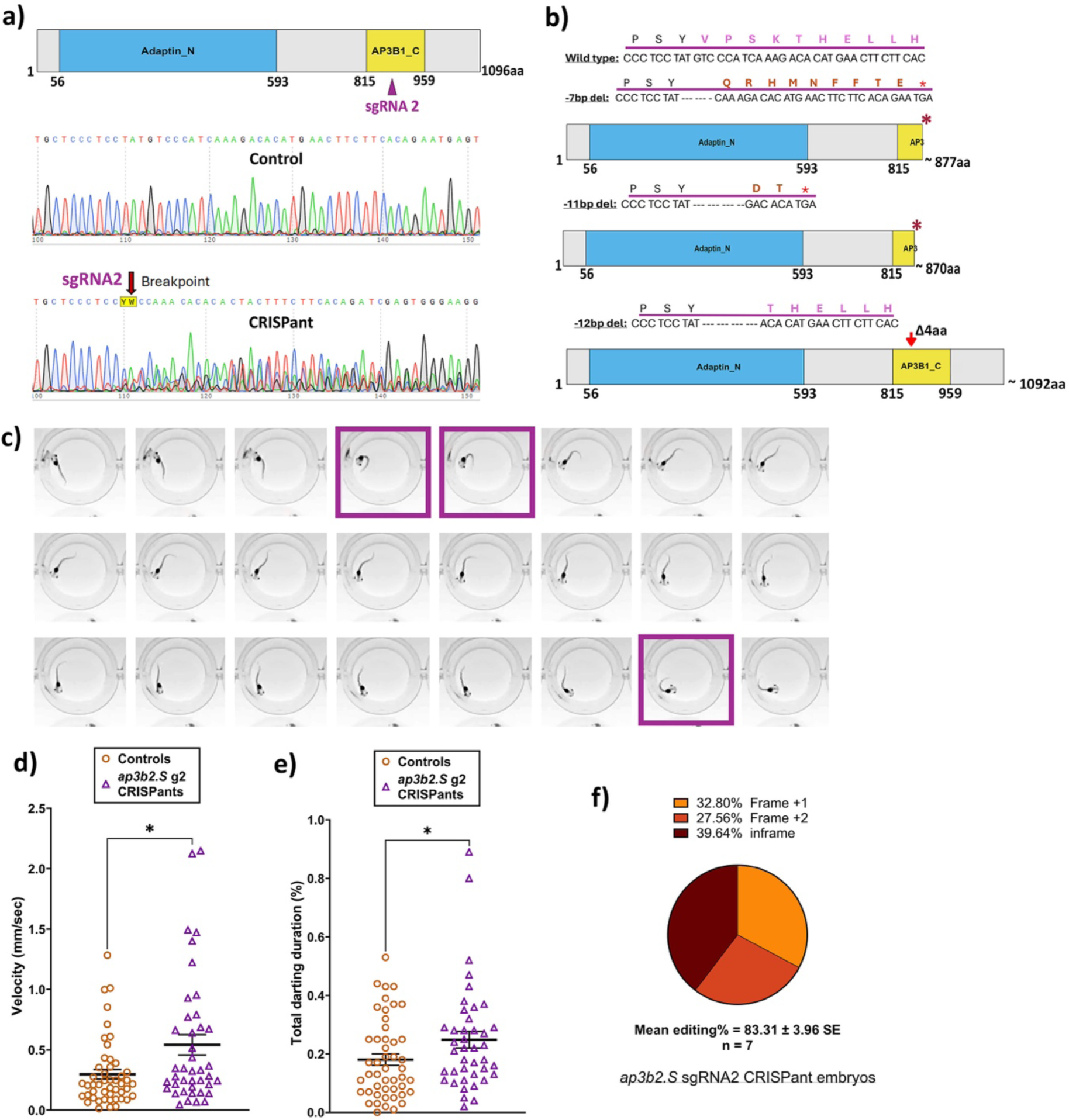
CRISPR/Cas9 disruption of *ap3b2* triggers premature protein truncation resulting in hyperactivity and seizure-like behaviour in CRISPants. (a) Schematic of the *Xenopus laevis* Ap3b2.S protein showing the conserved Adaptin_N and AP3B1_C domains. sgRNA2 target site is indicated, along with representative Sanger sequencing traces show degradation at the sgRNA2 cut site. (b) Predicted protein outcomes for the three most observed indels (-7, -11 and -12 bp deletions) generated by sgRNA2 (c) Representative frames from behavioural recordings at 50 fps of an *ap3b2* CRISPant tadpole. Frames highlighted in purple boxes show characteristic seizure-related C-shaped darting behaviour observed in CRISPants. (d-e) Comparison of (d) mean swim velocity (mm/s) and (e) time spent in darting behaviour (%) between *ap3b2* CRISPants (N = 41) compared with sibling controls (N = 48), unpaired t-test with Mann–Whitney, *P < 0.05. Horizontal bars indicate group means and error bars denote SEM. (f) Summary of CRISPR/Cas9 editing outcomes in 7 arbitrarily selected *ap3b2* CRISPant embryos from the same batch used in the behaviour experiments, confirmed by Sanger sequencing and TIDE analysis. Raw count data and statistical analysis can be found in Supplementary File 1.

### 3.2 Ca^2+^ imaging of the fore and midbrain of *ap3b2* CRISPant tadpoles reveals increased neural activity and increased synchrony between brain hemispheres

The marked behavioural hyperactivity and erratic swim episodes observed in *ap3b2* CRISPants prompted us to investigate whether Ap3b2 loss disrupts normal neuronal signalling during early brain development in our model. Using the same Ca²⁺ imaging protocol established in our DEE72 *neurod2* study (Banerjee et al., 2024), we recorded spontaneous *in vivo* activity from midbrain (MB) and forebrain (FB) regions in stage 47 tadpoles expressing GCaMP6s. CRISPants exhibited large, slow Ca²⁺ events that frequently persisted for more than two minutes, in contrast to the infrequent, low-amplitude fluctuations observed in controls (Figure 3a, <u>Supplementary video 2</u>).

**Figure 3.**
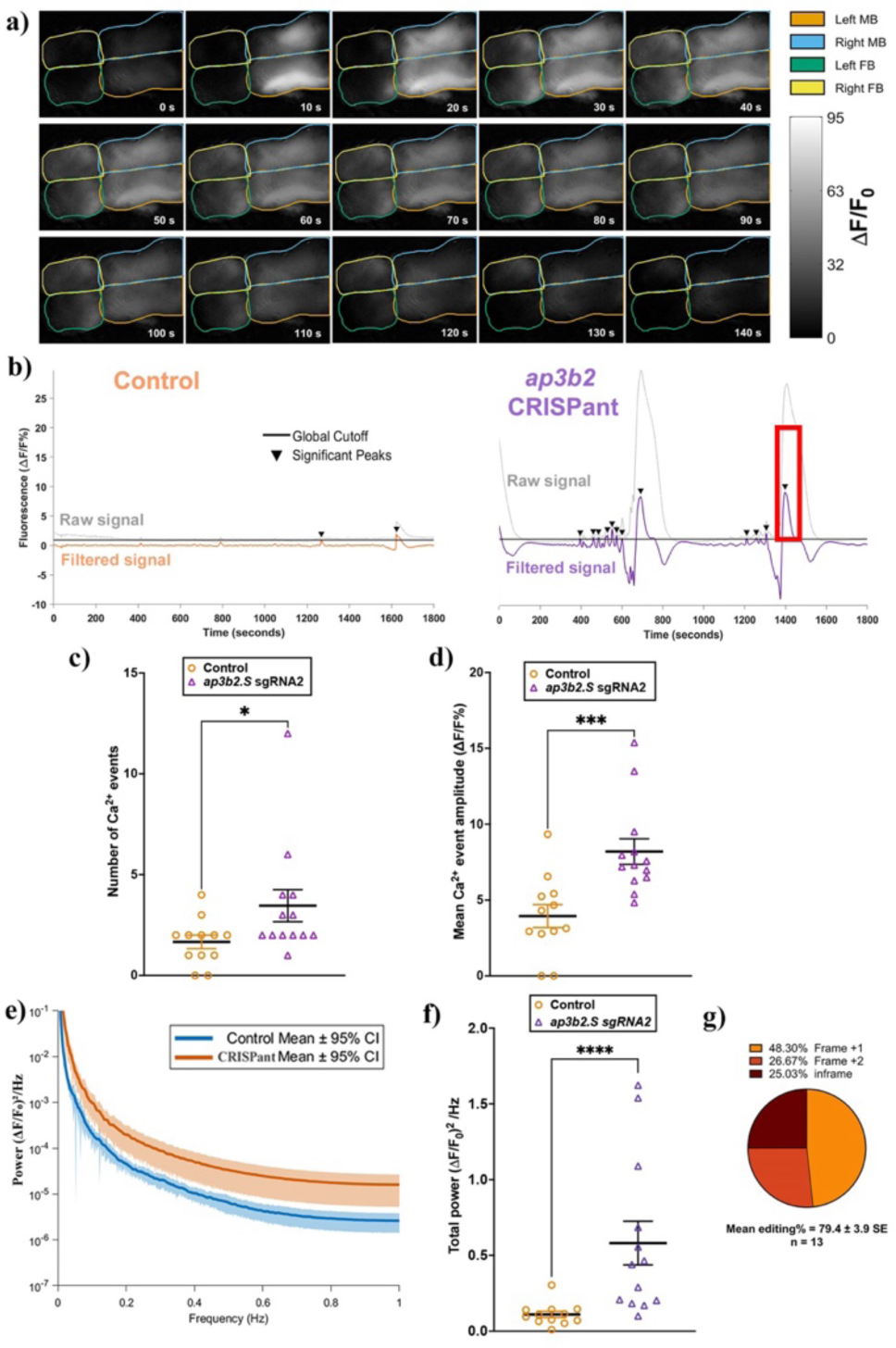
*Ap3b2* CRISPant brains have elevated spontaneous Ca^2+^ activity compared to unedited controls in CRISPants. (a) Representative still frames acquired every 10 s from *ap3b2* CRISPant tadpole brain during in vivo widefield Ca²⁺ imaging. GCaMP6s fluorescence intensity (lighter shades indicate higher ΔF/F_₀_%) is shown. Forebrain (FB) and midbrain (MB) hemispheres are outlined, the hindbrain lies outside the field of view. Time (s) is indicated in each frame. (b) Representative raw and high-pass-filtered ΔF/F₀% traces from a control tadpole (left) and an *ap3b2.S* CRISPant (right). Significant Ca^²⁺^ events were detected using a global threshold defined as 3× SD of all filtered control traces (black line); events exceeding this cutoff are marked by arrowheads. The red box indicates the time window in panel (a). (c,d) Comparison of Ca^²⁺^ event counts (c) and significant event amplitude (ΔF/F₀%) (d) in control (N = 12) and CRISPant tadpoles (N = 13).

To quantify this, ΔF/F₀ traces were first high-pass filtered at 0.005 Hz to remove baseline drift, and a global detection threshold, set at three times the SD of the filtered control cohort, was applied uniformly across all recordings (Figure 3b, Supplementary figure S6-S7). The brains of *ap3b2* CRISPant tadpoles showed a marked increase in spontaneous neural activity. The mean number of Ca²⁺ events was double that of controls (*ap3b2* CRISPants 3.46 +/- 0.80, unedited group 1.67 +/-0.33, p=0.028, Figure 3c). Mean event amplitudes were more than twice as high (*ap3b2* CRISPants 8.20 +/- 0.84, unedited group 3.94 +/- 0.76, p=0.0004, Figure 3d), indicating significantly more frequent and more intense events in CRISPants.

To assess whether loss of Ap3b2 alters the temporal organization of spontaneous Ca²⁺ activity in the tadpole brains, ΔF/F₀% signals recorded across fore and midbrain were analysed in the frequency domain using fast Fourier transform (FFT, Figure 3e). Consistent with prolonged and large-amplitude Ca²⁺ events observed, *ap3b2* CRISPant brains exhibited elevated low-frequency power across the 0.01–1 Hz range. To enable statistical comparison at the level of individual animals, integrated low-frequency spectral power (0.01–1 Hz) was calculated for each tadpole, (Figure 3f). Mean total spectral power was five times higher in *ap3b2.S* CRISPants compared with controls (CRISPants: 0.58 +/- 0.14; unedited group: 0.11 +/- 0.02; P < 0.0001), representing an approximately five-fold increase in low-frequency Ca²⁺ signal energy.

During qualitative inspection of the Ca^2+^ imaging data, spontaneous Ca²⁺ transients in *ap3b2* CRISPants appeared highly synchronous between hemispheres. To quantify this interhemispheric synchrony, we calculated Pearson correlation coefficients between left and right mid and forebrain centroid-based regions (Figure 4a). Control tadpoles displayed moderate left–right correlation within both mid- and forebrain regions (Figure 4b), consistent with normal developmental patterns of interhemispheric coordination. In *ap3b2* CRISPants, however, the correlation matrices showed uniformly elevated bilateral coupling across both brain regions (Figure 4c).

**Figure 4.**
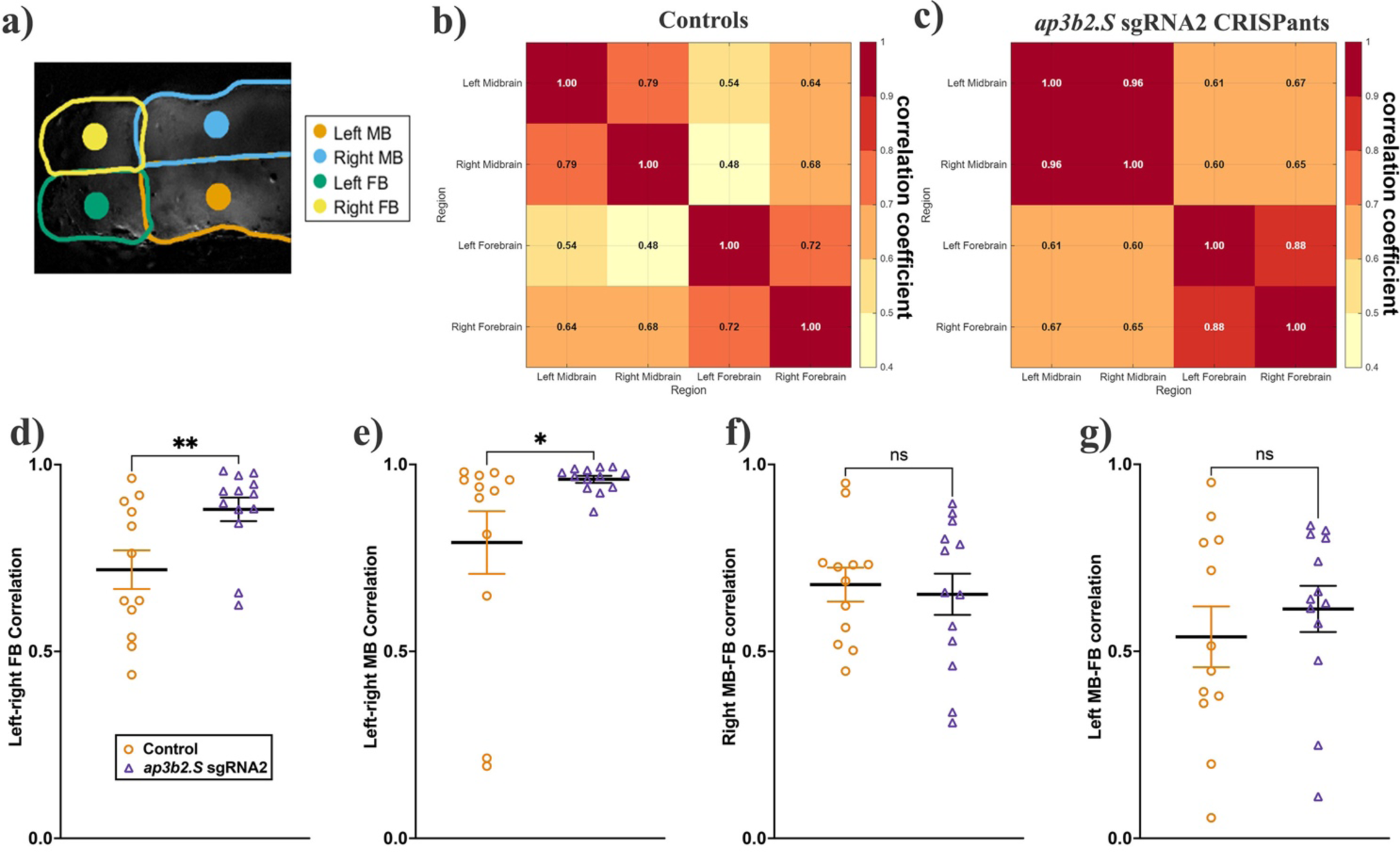
*Ap3b2* CRISPants exhibit increased interhemispheric synchrony in the midbrain and forebrain. (a) Schematic example illustrating the centroid-based regions of interest (ROIs) used for interregional correlation analysis. Fixed-area central ROIs (500 pixels each) were positioned at the geometric centroids of the left and right midbrain (MB), left and right forebrain (FB). (b, c) Group-averaged Pearson correlation coefficient matrices for unedited control tadpoles (b, N = 12) and *ap3b2* CRISPants (c, N = 13). (d–e) Comparison of brain regional synchrony, quantified as Pearson correlation coefficients, in controls and *ap3b2* CRISPants. Regions compared are (d) left-right MB and (e) left-right FB. Unpaired t-test with Mann Whitney, *P < 0.05, **P < 0.01. (f–g) Correlations between (f) right MB-FB, and (g) left MB-FB, unpaired t-tests with Mann Whitney, ns, not significant. Raw correlation values and full statistical details are provided in Supplementary File 1.

Quantitative analysis showed that mean left–right midbrain synchrony was significantly higher in *ap3b2* CRISPants (0.96 +/- 0.01, N = 13 tadpoles) than in unedited controls (0.79 +/- 0.08, N = 12) (p = 0.04; Figure 4d). A similar increase was observed in the forebrain, where mean left–right forebrain synchrony was also significantly elevated in CRISPants (0.88 +/- 0.03) compared with controls (0.72 +/- 0.05) (p = 0.01; Figure 4e). By contrast, heterotopic correlations did not differ significantly between *ap3b2* CRISPants and controls (right midbrain-forebrain, Figure 4f, p=0.72, left midbrain-forebrain, Figure 4g, p=0.47). Together, the observed increased Ca^2+^ activity and enhanced interhemispheric synchrony in *ap3b2* CRISPant brains are consistent with a highly synchronised network state, which could underlie the seizure susceptibility of both CRISPant tadpoles and DEE48 patients.

Individual data points represent single tadpoles; horizontal bars denote group means and error bars indicate SEM. Groups were compared using unpaired t-test with Mann Whitney, *P < 0.05, ***P < 0.001. (e) FFT-derived power spectral densities of whole-brain Ca²⁺ signals for control and *ap3b2* CRISPant tadpoles. Solid lines represent group means and shaded envelopes indicate 95% confidence intervals. (f) Integrated low-frequency spectral power (0.01–1 Hz; (ΔF/F₀)²/Hz, calculated as the area under the power spectrum for each tadpole. Data are from control (N = 12) and *ap3b2.S* CRISPant tadpoles (N = 13). Individual data points represent single tadpoles; horizontal bars denote group means and error bars indicate SEM. Groups were compared using unpaired t-test with Mann–Whitney correction, ***P < 0.001, ****P < 0.0001. (g) Summary of CRISPR/Cas9 editing outcomes in the CRISPant group, determined by Sanger sequencing and TIDE analysis. Raw count data and full statistical details are provided in Supplementary File 1.

### 3.3 *Ap3b2* CRISPant brains have reduced expression of genes associated with monovalent cation transport, BBB function, inhibitory GABA neurotransmission and axon guidance

The pronounced changes in brain activity prompted us to examine whether *Ap3b2* loss is accompanied by transcriptional changes in pathways governing neural circuit function and homeostasis. Prior developmental transcriptome mapping in *X. laevis* tadpole brain development showed both region and stage-specific shifts in neuronal, progenitor, and synaptic gene programmes, supporting the suitability of this system for detecting mutation-induced network changes (Ta et al., 2021). To determine what developmental changes predispose DEE48 brains to seizure activity, we performed RNA-seq on pooled stage 47 unedited control and *ap3b2.S* CRISPant brains, the same stage at which behavioural and neural activity assays were conducted.

Principal component analysis (PCA) showed that genotype was the strongest driver of gene expression in our dataset, indicating an effect of *ap3b2* loss on the developing brain transcriptome (Figure 5a). EdgeR analysis was used to identify differentially expressed genes (DEG: FDR < 0.05, log₂FC > 1) revealed a strongly asymmetric response: 1,078 genes were significantly downregulated in CRISPant brains compared to control unedited brains, whereas only 28 were upregulated, with median fold changes of ∼2.3-fold decrease and ∼2.1-fold increase, respectively. The volcano plot illustrates this bias towards downregulation, with top-ranked genes selected using a Euclidean distance-based ranking that integrates both effect size (log₂ FC) and statistical significance (–log₁₀ adjusted P value). Top ranked genes included the endothelial junction gene *jcad.S*, the transporter *slc16a2.L*, and transcriptional regulators *bach2.L* and *zbtb20.S*, while the most upregulated genes (e.g., *ikbke.S, cfi.L*, *cps1.S*, *mpc2.L*, *krt12.5.L*, *krt12.5.S*, *krt62.S*) are associated with immune, metabolic and cytoskeletal functions (Figure 5b).

**Figure 5.**
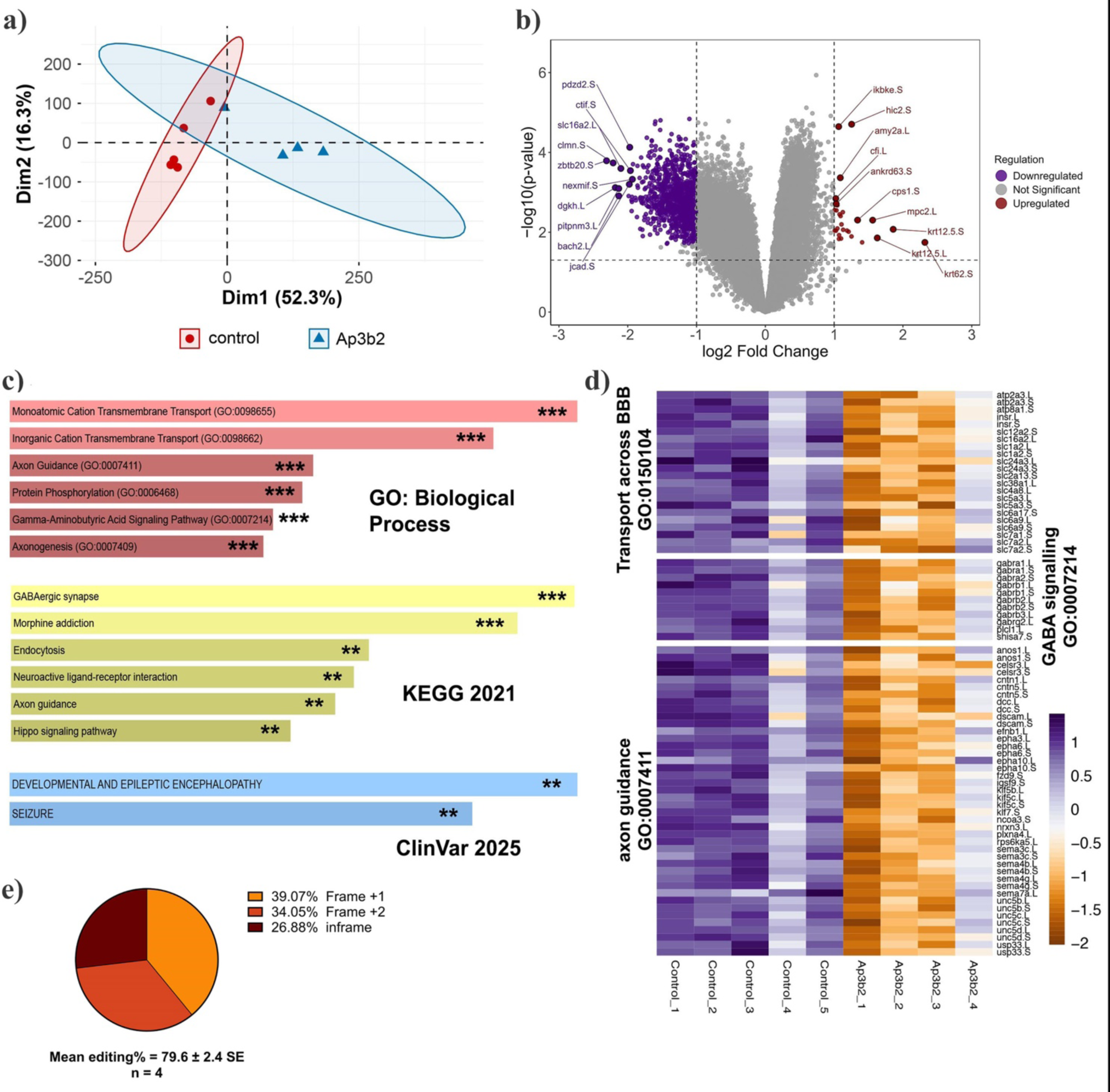
*ap3b2* disruption downregulates BBB transport, GABA signalling, and axon guidance pathways in CRISPant tadpole brains. a) PCA plot of normalized read counts of the five control samples and four *ap3b2* **^⁻/⁻^** (mosaic) samples, each sample is derived from six pooled brains. b) Volcano plot of EdgeR differentially expressed genes, (DEG) with FDR threshold of <0.05 and Log2 fold change of >1. The 10 most significant up and down regulated genes are labelled. c) Selected overrepresented ontologies calculated with EnrichR for down regulated DEG, the top 6 hits are shown for GO: biological process and KEGG_2021, for ClinVar the only two significant hits are shown (PAdj <0.05). d) Heatmap of z-scores for three down overrepresented ontologies, down regulated in CRISPants. PAdj ** <0.01, *** <0.001. e) Summary of tadpole editing in the *ap3b2* CRISPant group, confirmed by Sanger sequencing and TIDE analysis. Supplementary data for (c) and (e), as well as the custom brain background gene list, can be found in Supplementary file 1.

The upregulated DEG group was too small to generate any significantly enriched pathway data. Functional enrichment analysis of the downregulated set was conducted against a background list of stage matched brain-expressed genes, using Enrichr (Chen et al., 2013, Kuleshov et al., 2016, Xie et al., 2021). Many pathways and processes were found to be significantly overrepresented in this *ap3b2* CRISPant down-regulated gene set, including ion transport, inhibitory signalling, blood brain barrier formation, and establishment of neural connectivity (axonogenesis/axon guidance). Both GO biological process and KEGG pathway analyses highlighted monovalent and inorganic cation transport, GABAergic synapse/signalling, axon guidance, endocytosis, and neuroactive ligand-receptor interaction. Two ClinVar terms “Developmental and Epileptic Encephalopathy” and “Seizure” were significantly overrepresented (Figure 5c,d). Genes associated with transport across the BBB (GO:0150104), included BBB-associated transporters and endothelial genes: *slc16a2.L*, *slc11a2.L*, *slc11a2.S*, *abcg1.L*, *abcg1.S*, *cldn11.S*, *flt1.L*, *kdrl.L*). Down regulated genes were also enriched for GABA signalling (GO:0007214), encompassing multiple GABA_A receptor subunits and synthetic machinery (*gabra1.L*, *gabra2.L*, *gabrb1.L*, *gabrb1.S*, *gabrb2.L*, *gabrb2.S*, *gabrb3.L*, *gabrg2.L*, *gad2.L*); and axon guidance (GO:0007411), including guidance receptors and ligands such as *plxna4.L*, *sema3c.L*, *sema3c.S*, *sema4b.S*, *sema4g.L*, *sema4g.S*, *sema7a.L*, *unc5b.L*, *unc5b.S*, *unc5c.L*, *unc5c.S*, *unc5d.S* (Figure 5d). These data indicate that *ap3b2* knockdown leads to a broad, pathway-level downregulation of BBB transport, inhibitory neurotransmission, and axon wiring programmes. High mean editing efficiency in the CRISPant cohort (∼80%; Figure 5e) supports a direct link between AP-3 disruption and these transcriptomic changes.

### 3.4 Increased early BBB permeability in *ap3b2* CRISPants

In our previous *neurod2* CRISPant DEE72 model, we demonstrated that early-onset seizures coincide with pronounced BBB leakage, with rapid sodium fluorescein (NaF) dye escape occurring despite otherwise normal brain morphology. Notably, short-term losartan treatment reduced both dye leakage and seizure burden, supporting a mechanistic link between BBB permeability and epileptogenesis rather than a secondary consequence of seizures (Banerjee et al., 2024). Our analysis of down regulated genes in the brains of *ap3b2* CRISPant tadpoles (Figure 5d) showed significant enrichment of pathways related to BBB transport suggesting impaired or developmentally delayed barrier function. This raised the possibility that *ap3b2* loss, similar to *neurod2* haploinsufficiency, compromises BBB integrity.

We therefore assessed whether *ap3b2* CRISPants have altered BBB integrity, by monitoring the diffusion of intraventricularly injected NaF across four timepoints (2, 5, 10, and 20 minutes post-injection). Visual inspection of CRISPant tadpoles revealed no gross abnormalities in overall brain morphology compared with controls, consistent with observations previously reported in the DEE72 tadpole CRISPant model (Banerjee et al., 2024). As expected, control tadpoles showed a slow and progressive spread of fluorescence beyond the ventricular space, consistent with low-level physiological leakage from a normally developing BBB. In contrast, *ap3b2* CRISPants displayed markedly accelerated dye escape, with significantly higher fluorescence outside the brain and in kidneys, apparent after 2 minutes (P= 0.0212, Figure 6c), indicating an immediate increase in BBB permeability. Although the difference between groups partially converged at 5–10 minutes, reflecting rapid equilibration, once the barrier is breached the later phase of the assay revealed a clear divergence. By 20 minutes, most fluorescein had already cleared from both the brain and surrounding tissue in CRISPants, whereas controls continued to show a gradual outward leak and retained a visible dye reservoir (Figure 6c, Supplementary figure S8). Together, these data demonstrate that loss of approx. 80% of ap3b2, resulting from editing (Figure 6d) causes early-onset and transiently heightened BBB permeability, which could represent an early pathological feature of AP3B2-associated DEE.

**Figure 6.**
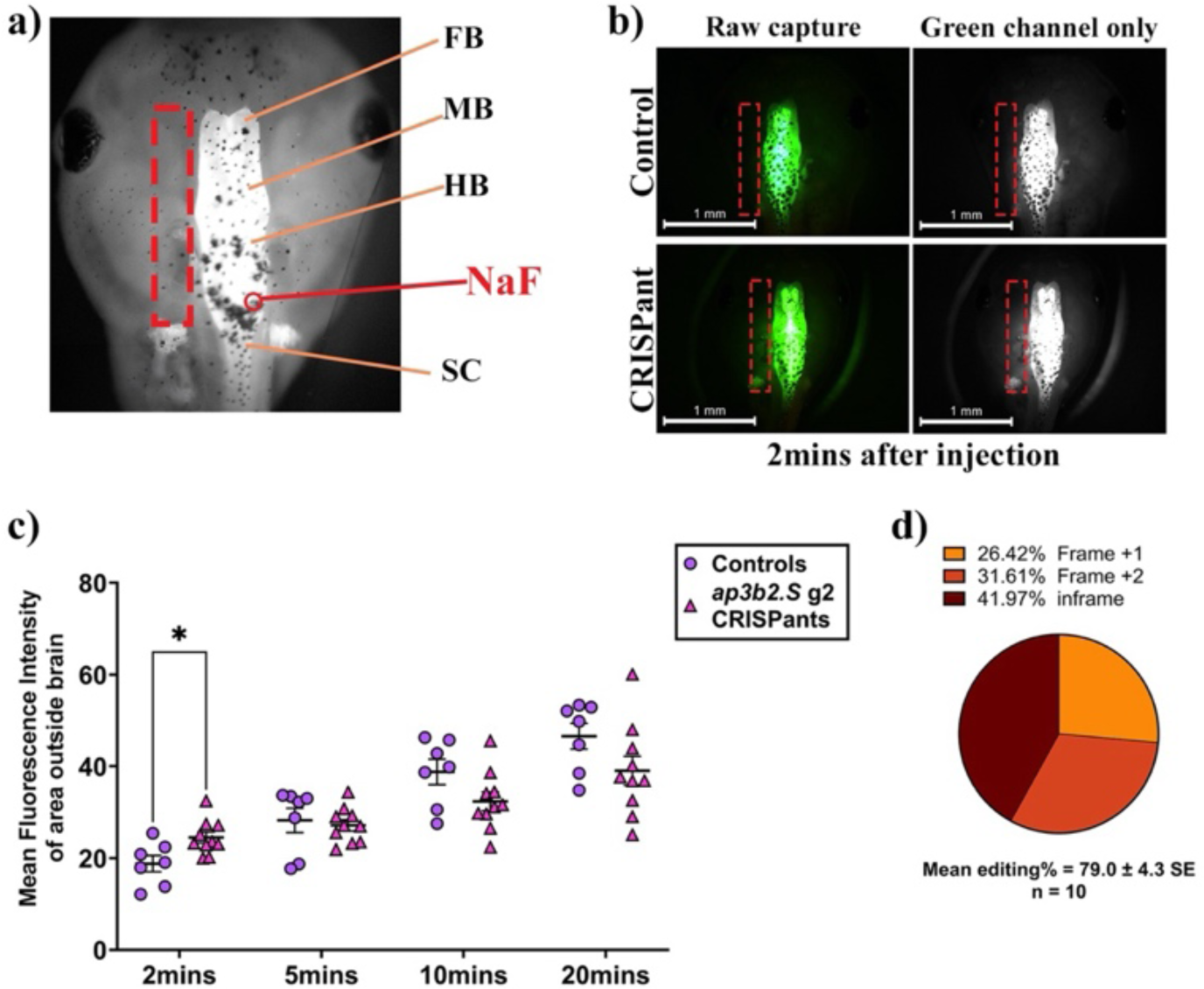
Rapid early sodium fluorescein dye leakage in *ap3b2* CRISPants reveals a markedly compromised BBB. (a) Schematic dorsal view of an NF stage 47 *Xenopus laevis* tadpole head showing forebrain (FB), midbrain (MB), hindbrain (HB), and spinal cord (SC), and the site of sodium fluorescein (NaF) microinjection into the 4th ventricle (red arrow). The dashed rectangle indicates the ROI outside the brain from which fluorescence intensity was quantified. b) Representative images of NaF-injected controls (top row) and *ap3b2.S* CRISPants (bottom row) 2 minutes after microinjection (raw capture and green channel only). (c) Plot of mean fluorescence intensity (MFI) detected outside tadpole brain at 2, 5, 10 and 20 minutes post NaF injection in CRISPant tadpoles (N = 10) compared to controls (N = 7), Repeated measures 2-way ANOVA with Tukey’s multiple comparisons test, *P < 0.05. (d) Summary of CRISPR/Cas9 editing outcomes in the CRISPant group, determined by Sanger sequencing and TIDE analysis. Raw count data and full statistical details are provided in Supplementary File 1.

### 3.5 Losartan treatment reduces mean swimming velocity, without significantly altering Ca^2+^ dynamics of *ap3b2* CRISPant brains

Drug-refractory seizures remain a major clinical challenge in developmental and epileptic encephalopathies (DEEs), with many patients showing limited or no response to conventional anti-seizure medications. This has prompted increasing interest in repurposed therapeutics that act on non–ion-channel pathways, particularly agents with established clinical safety profiles and anti-inflammatory properties. One such candidate is losartan, a widely used angiotensin II type-1 receptor antagonist that modulates neuroinflammatory signalling by increasing thrombospondin-1 (TSP1) expression and thereby regulating latent TGF-β activation (Figure 7a). Losartan has been shown to suppress the development of chronic seizures in rodent models of traumatic brain injury and, more recently, to acutely reduce seizure burden in the *X. laevis neurod2* CRISPant model (Banerjee et al., 2024). These prior findings implicate TGF-β–associated inflammatory and blood–brain barrier–linked pathways as contributors to seizure susceptibility and raise the possibility that similar mechanisms contribute to the pathology of other DEEs, such as DEE48.

**Figure 7.**
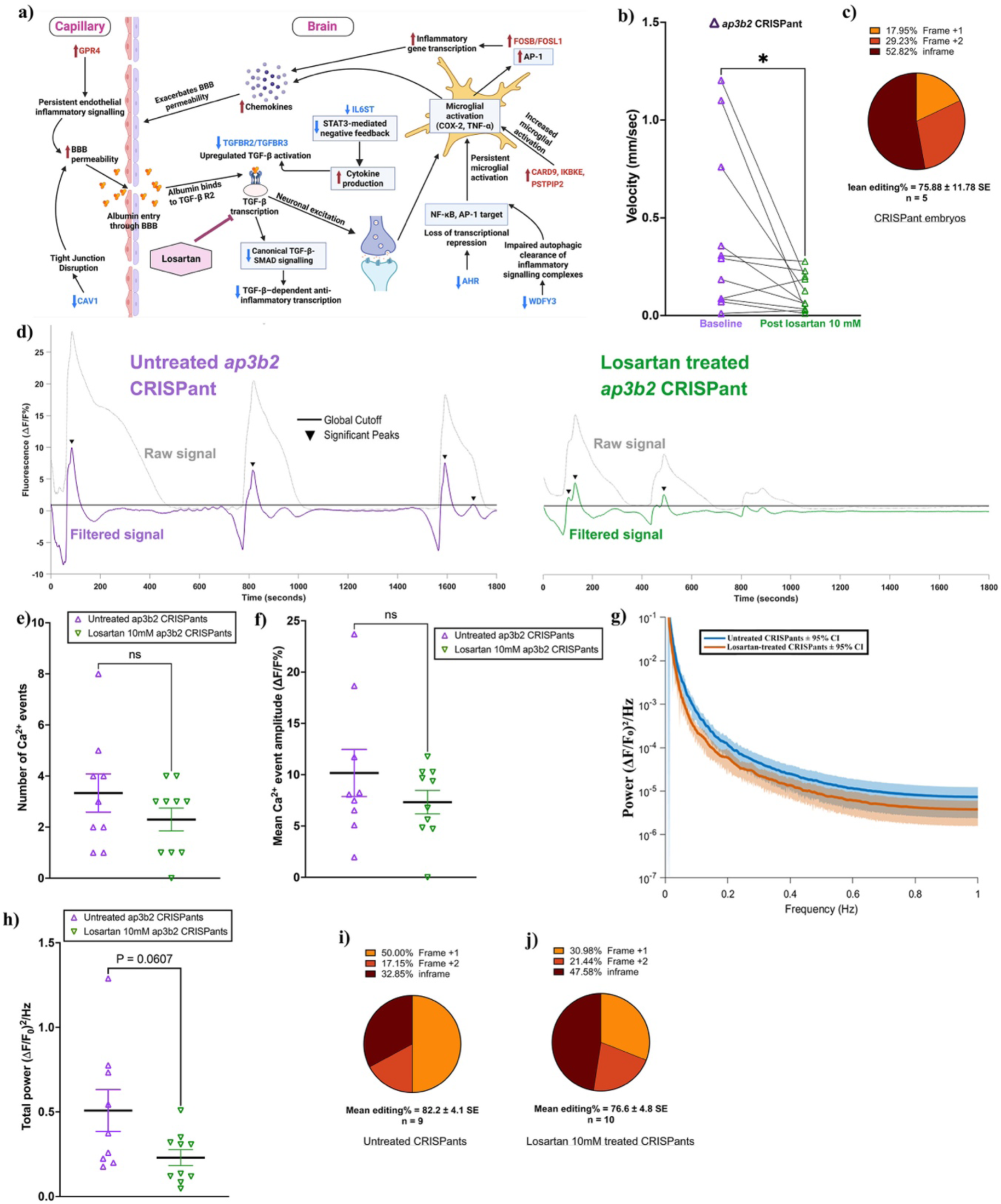
Losartan treatment reduces aberrant calcium activity and hyperactivity in *ap3b2* CRISPants. (a) Conceptual schematic of *ap3b2* CRISPant differentially expressed genes associated with loss of anti-inflammatory regulatory control, BBB dysfunction, and neuronal hyperexcitability relevant to TGF-β receptor–SMAD signalling, indicating a hypothetical mode of action for losartan. (Created in BioRender). (b) Paired comparison of mean swim velocity (mm/s, over 1 hour) in *ap3b2* CRISPant tadpoles before and after 10 mM losartan treatment. Lines connect individual tadpoles, and the effect of losartan was tested using a Wilcoxon paired t-test, *P < 0.05. (c) Summary of CRISPR/Cas9 editing outcomes for embryos used in the losartan phenotype experiments, determined by Sanger sequencing and TIDE analysis. (d) Representative whole-brain Ca^2+^ signals (raw and filtered ΔF/F₀ traces) showing activity from untreated *ap3b2.S* CRISPants (left) and losartan-treated CRISPants (1 hour, 10 mM, right). Black lines indicate the global event detection threshold (3× control SD), with arrowheads marking significant Ca²⁺ events. (e,f) Comparison of the numbers of Ca^²⁺^ events (e) and mean event amplitude (f) detected in untreated (N = 9) and losartan treated (10 mM; N = 10) *ap3b2* CRISPant tadpoles. Groups were compared using unpaired Welch’s t-tests, ns = not significant (P > 0.05). (g) FFT-derived power spectral densities of spontaneous whole-brain Ca²⁺ activity in untreated and losartan-treated *ap3b2* CRISPants, plotted on a logarithmic scale. Solid lines indicate group means and shaded envelopes represent 95% confidence intervals across animals. (h) Comparison of integrated low-frequency spectral power density (0.01–1 Hz; (ΔF/F_₀_)^²^/Hz) between untreated (N = 9) and losartan-treated (10 mM; N = 10) *ap3b2* CRISPants. Individual data points on scatterplots represent single CRISPant tadpoles; horizontal bars denote group means and error bars indicate SEM. Statistical significance was assessed an unpaired Welch’s t-test, with the exact p value shown. (i,j) Distribution of CRISPR editing outcomes in untreated (i) and losartan-treated (j) CRISPant tadpoles, quantified by Sanger sequencing and TIDE analysis. Raw data and full statistical analyses are provided in Supplementary File 1.

To see if there was evidence for this in our *ap3b2* CRISPR brain transcriptomes, we looked at several key anti-inflammatory regulators, including *tgfbr2*, *tgfbr3*, *il6st*, *ahr*, *wdfy3*, and *cav1*, each of which normally constrains CNS inflammatory signalling (Table 2). Loss of TGFBR2/3 weakens canonical TGF-β–SMAD signalling, a pathway required to maintain microglia and astrocytes in a homeostatic, non-reactive state (Zöller et al., 2018, Luo, 2022, Blair et al., 2011, Duesman et al., 2023). Reduced IL6ST further diminishes STAT3-mediated cytokine negative feedback (Murakami et al., 2019, Rose-John, 2018, Cekanaviciute and Buckwalter, 2016), while decreased AHR and WDFY3 remove important transcriptional and autophagic brakes on glial activation (Wheeler et al., 2017, Wang et al., 2023, Filimonenko et al., 2010, Fox et al., 2020). Concurrent downregulation of *cav1* suggests impaired BBB stability, increasing susceptibility to peripheral inflammatory mediators (Huang et al., 2018, Trevino et al., 2024).

**Table 2.** Neuroinflammation-associated DEG downregulated in *ap3b2* CRISPant tadpole brains.

| *X. laevis* gene | Human ortholog | Log<sub>2</sub>FC | PValue | FDR |
| --- | --- | --- | --- | --- |
| *il6st.L* | *IL6ST* | -1.0822 | 0.0003 | 0.0098 |
| *tgfbr2.L* | *TGFBR2* | -1.6293 | 0.0018 | 0.0133 |
| *tgfbr2.S* |  | -1.4140 | 0.0002 | 0.0098 |
| *tgfbr3.L* | *TGFBR3* | -1.0863 | 0.0056 | 0.0224 |
| *tgfbr3.S* |  | -1.7020 | 0.0046 | 0.0203 |
| *ahr.L* | *AHR* | -1.6638 | 0.0062 | 0.0236 |
| *ahr.S* |  | -1.4367 | 0.0011 | 0.0115 |
| *wdfy3.S* | *WDFY3* | -1.1333 | 0.0007 | 0.0105 |
| *cav1.L* | *CAV1* | -1.2217 | 0.0014 | 0.0123 |

In parallel, several potent pro-inflammatory genes were upregulated (Table 3), including *card9*, *ikbke*, *pstpip2*, *fosb*, *fosl1*, *gpr4* and *cfi*, which respectively activate NF-κB, interferon, AP-1, complement and endothelial inflammatory pathways (Hara et al., 2007, Zhong et al., 2018, Verhelst et al., 2013, Clément et al., 2008, Cassel et al., 2014, He et al., 2022, Dong et al., 2013, Gomez-Arboledas et al., 2021). Together, this pattern reflects a shift from protective, TGF-β–dependent immune homeostasis toward a state of heightened innate immune activation. Thus, downregulation of TGF-β signalling components, combined with induction of cytokine- and danger-associated transcripts, provides a mechanistic framework by which disrupted Ap3b2 function may predispose the developing brain to persistent, TGF-β–associated neuroinflammation (Figure 7a).

**Table 3.** Neuroinflammation-associated genes upregulated in *ap3b2* CRISPant tadpole brains.

| *X. laevis* gene | Human ortholog | Log<sub>2</sub>FC | PValue | FDR |
| --- | --- | --- | --- | --- |
| *card9.L* | *CARD9* | 1.1067 | 0.0064 | 0.0239 |
| *ikbke.S* | *IKBKE* | 1.0658 | 0.0000 | 0.0098 |
| *pstpip2.S* | *PSTPIP2* | 1.1003 | 0.0144 | 0.0401 |
| *fosb.L* | *FOSB* | 1.2522 | 0.0127 | 0.0369 |
| *fosl1.S* | *FOSL1* | 1.1257 | 0.0031 | 0.0168 |
| *gpr4.L* | *GPR4* | 1.2102 | 0.0097 | 0.0307 |
| *cfi.L* | *CFI* | 1.0237 | 0.0014 | 0.0124 |

To test whether targeting TGF-β associated inflammatory signalling could mitigate the seizure phenotype observed in *ap3b2* CRISPants, tadpoles were treated with 10 mM losartan and assessed using behavioural and Ca²⁺ imaging assays (**Error! Reference source not found.**). The addition of individual paired measurements demonstrated decreased swim velocity of 9/11 *ap3b2* CRISPant tadpoles following a 1 hour treatment with 10 mM losartan (Figure 7b, mean velocity 0.41+/-0.13 mm/sec before treatment and 0.11+/-0.03 mm/sec after treatment, p=0.02). Embryo sequencing from this batch of CRISPants confirmed high editing efficiency (∼76%). For assessment of Ca^2+^ signalling, it was not possible to use the same tadpoles, so *ap3b2* CRISPants were arbitrarily assigned to treatment (10 tadpoles) or control (9 tadpoles) groups.

GCaMP6S live Ca^2+^ imaging appeared to show a partial suppression of abnormal activity in the losartan treated group (Figure 7d, <u>Supplementary video 3</u>, Supplementary figures S9 and S10). Further analysis of the Ca^2+^ imaging data confirmed that untreated *ap3b2* CRISPant brains discharged frequent, high amplitude Ca²⁺ events, as previously shown. While the losartan treated group generated 31% fewer such events (control mean 3.33 ± 0.87; treated mean 2.30 +/- 0.73), this did not represent a significant reduction (P = 0.256; Figure 7e). Similarly, although the amplitudes of the losartan treated group Ca^2+^ events (ΔF/F₀%) were on average 28% lower than in untreated *ap3b2* CRISPants (control mean 10.17 +/- 2.56; treated mean 7.32 +/- 2.14), the effect was not significant (P=0.289, Figure 7f). Group-averaged power spectral density curves (Figure 7g, 0.01-1 Hz), indicated that losartan-treated *ap3b2* CRISPants tend to show less slow calcium fluctuations. Total spectral power was 60% lower in the treated group (control mean 0.508 +/- 0.124; losartan treated mean 0.230 +/- 0.047; P = 0.061; Figure 7h). Analysis of CRISPR/Cas9 editing outcomes revealed no difference in editing efficiencies and indel distributions between untreated and losartan-treated *ap3b2* CRISPants (Mann-Whitney test, p=0.39, Figure 7i,j; Supplementary figure S11). The observed significant reduction in swimming velocity following losartan treatment of individual tadpoles, together with a trend of decreased seizure-like brain activity, measured through live Ca^2+^ monitoring of equivalent *ap3b2* CRISPant groups, suggests that targeting neuroinflammatory pathways may have beneficial effects in reducing seizure activity in DEE48.

## 4 Discussion

### 4.1 Ap3b2 loss of function generates a robust DEE48-like phenotype in *X. laevis* tadpoles

We set out to mimic the homozygous loss of function variants found in human patients with DEE48, using F_0_ CRISPant tadpoles. While these tadpoles are mosaic, greater than 80% of *ap3b2.S* genes were found to be edited on average. Two thirds of edits are predicted to result in a truncating frameshifts, seen in some patients, and the remainder correspond to a 12 bp in-frame deletion, of unknown consequence, but located in the conserved AP3B1-C terminal domain. Because *AP3B2*-associated DEE48 is an autosomal-recessive disorder, the observed robust phenocopy in tadpoles strongly suggests that loss of these four amino acids is also pathogenic. Disruption of the *X. laevis ap3b2* homeologue produced a behavioural and neurophysiological phenotype that parallels the clinical presentation of DEE48. *ap3b2* CRISPant tadpoles exhibited increased mean swimming velocity, more time spent darting, behavioural signatures strongly reminiscent of the seizure-like motor episodes reported in individuals with biallelic AP3B2 loss-of-function variants (Alizadeh et al., 2025, Dilber et al., 2022, Assoum et al., 2016, Anazi et al., 2017). Seizures and hyperactivity were also reported in an *ap3b2^-/-^* mouse model (Nakatsu et al., 2004), suggesting functional conservation at least across vertebrates. Ca²⁺ imaging of brain activity revealed increased occurrence of spontaneous, large-amplitude, prolonged Ca^2+^ transients and elevated interhemispheric synchrony consistent with network hyperexcitability characteristic of epileptic encephalopathy.

AP3B2 encodes the neuron-specific β-subunit of the adaptor protein-3 (AP-3) complex, required for synaptic vesicle and endolysosomal cargo trafficking. In mice, selective loss of neuronal AP-3B alone is sufficient to cause spontaneous seizures and increased seizure susceptibility in the absence of gross brain malformations, driven by disturbed synaptic vesicle function (Nakatsu et al., 2004). Together with human genetic evidence, these findings have established defective synaptic vesicle trafficking as a core pathogenic mechanism underlying AP3B2-associated DEE. Consistent with this model, increased spontaneous brain activity and interhemispheric synchrony observed in *ap3b2* CRISPant tadpole brains likely arises from a primary synaptic trafficking defect that disrupts inhibitory–excitatory balance during critical periods of neural circuit assembly.

Our results also align with a growing body of work demonstrating the translational power of rapid *in vivo* CRISPR-based modelling of rare genetic epilepsies in aquatic vertebrates. CRISPR targeted *spout1* disruption in zebrafish resulted in epileptiform activity and neurodevelopmental abnormalities akin to those found in patients with compound heterozygous mutations in SPOUT1, confirming pathogenicity (Liu et al., 2024). In the *neuroD2* (DEE72) haploinsufficient CRISPant model, spontaneous seizure-like behaviour and increased neural activity were observed despite preserved gross brain morphology (Banerjee et al., 2024), a pattern strikingly recapitulated in the *ap3b2*^-/-^(mosaic) CRISPants. The *ap3b2.S* CRISPant model therefore joins a growing class of vertebrate DEE systems in which targeted gene perturbation directly yields a reproducible encephalopathic phenotype, strengthening genotype–phenotype interpretation for rare variants. The early accessibility of both zebrafish and *Xenopus* CRISPant models highlights these model organisms as powerful platforms for interrogating early disease mechanisms and evaluating candidate modifiers in DEE.

### 4.2 *Ap3b2* loss reveals early blood–brain barrier fragility and suggests altered neuroinflammation

Early BBB dysfunction is increasingly recognised as a key contributor to seizure susceptibility and epileptogenesis. Increased BBB permeability permits serum proteins, ions, and inflammatory mediators to enter the brain parenchyma, disrupting astrocytic regulation of potassium and glutamate homeostasis and promoting network hyperexcitability (Swissa et al., 2019). In paediatric epilepsies, BBB instability correlates with higher seizure burden and drug resistance, supporting the view that barrier dysfunction is a driver of disease severity rather than a secondary consequence (Kimizu et al., 2018). Because the developing brain is particularly sensitive to neurovascular disruption, even transient BBB opening may have lasting effects on circuit maturation and seizure risk (Moretti et al., 2015).

A striking phenotype in *ap3b2* CRISPants was the rapid and pronounced leakage of sodium fluorescein from the ventricular system, compared to unedited siblings, indicating early and severe BBB compromise. This physiological defect was strongly supported by transcriptomic data showing coordinated downregulation of endothelial solute carriers, transporters, adhesion molecules, and tight-junction–associated genes. The rapid leakage followed by accelerated washout is consistent with early, high-velocity barrier failure rather than gradual diffusion, suggesting that BBB integrity is compromised almost immediately upon dye entry. Furthermore, transcriptome analysis of *ap3b2* CRISPant brains found that genes associated with the BBB were overrepresented in the set of down-regulated differentially expressed genes. These genes included solute carriers, endothelial transporters, adhesion molecules, and tight-junction components such as cldn11. Similar patterns are characteristic of epileptogenic tissue and experimental seizure models (Chen et al., 2020, Salman et al., 2017). Consistent with these molecular changes, *ap3b2* CRISPants exhibited rapid sodium fluorescein leakage despite grossly normal brain morphology, indicating impaired barrier integrity. Together, these data suggest that AP3B2 loss disrupts neurovascular maturation, creating a permissive environment for aberrant extracellular signalling and reduced homeostatic control. A similarly less robust BBB was demonstrated in our recent model of DEE72 (Banerjee et al., 2024), suggesting a common mechanism leads to this phenotype. It is not yet known whether BBB leakiness is a cause or effect of seizure activity, but this commonality points towards the latter.

Currently is it not known whether human DEE patients also have a compromised BBB, but the broader epilepsy literature strongly supports this interpretation. Across neonatal hypoxic–ischaemic injury, traumatic brain injury, and drug resistant epilepsy, early BBB opening permits albumin and cytokine entry into the brain, triggering astrocytic activation, impaired ion buffering, and downstream network hyperexcitability (Dadas and Janigro, 2019, Goasdoue et al., 2019, Specchio et al., 2010). Albumin-driven activation of TGF-β signalling in astrocytes is a well-established ictogenic mechanism, and blockade of this pathway prevents epileptogenesis in multiple experimental models (Gorter et al., 2015, Librizzi et al., 2012).

In human patients with DEE, infantile spasms are often among the first seizure types detected. It has long been the custom to treat infantile spasms with ACTH or steroid therapy, which offers a drastic but often effective solution to neuroinflammation, and the resulting damage to developing brains, that comes with unrelenting seizure activity. More generally, the role of neuroinflammation in epilepsy is emerging as an important but often overlooked potential therapy target (Sanz et al., 2024). While we did not find neuroinflammatory pathways to be overrepresented in our transcriptome analyses of *ap3b2* CRISPant brains, *ikbke.S* was one of the most significantly up-regulated genes, prompting us to look for other DEG associated with neuroinflammation (Tables 1 and 2). Additionally, *ap3b2* CRISPants brains showed significant transcriptomic suppression of multiple components associated with TGF-β responsiveness, including of *tgfbr2, tgfbr3, il6st* and *skil.* Although inflammatory activation was not directly measured, these pathways are repeatedly implicated in seizure susceptibility, BBB stability, and neurovascular regulation (Chen et al., 2020, Okamoto et al., 2010). The convergence of reduced TGF-β–associated signalling, BBB dysfunction, and network hyperexcitability suggests a coordinated failure of neuronal, glial, and vascular regulatory systems rather than isolated defects.

The convergence of BBB leakage, suppression of neurovascular regulatory genes, and increased and hypersynchronous neural activity supports a model in which AP3B2 loss creates a permissive environment for persistent hyperexcitability during development. The shared neurovascular phenotype observed across DEE48 and DEE72 *Xenopus* models further highlights BBB fragility as a conserved mechanistic vulnerability in genetically driven epileptic encephalopathies, with important implications for understanding disease severity and therapeutic responsiveness.

### 4.3 Coordinated transcriptomic changes links molecular, behavioural, and Ca^2+^ imaging phenotypes

Transcriptomic studies across human epilepsy and experimental seizure models consistently show that epileptic encephalopathies arise from broad, coordinated disruption of neuronal, glial, immune, and vascular gene networks rather than isolated pathway defects. Large-scale RNA sequencing of epileptic tissue reveals widespread suppression of neuronal and synaptic programmes alongside changes in immune and vascular signalling (Guelfi et al., 2019, Iacobaş and Velíšek, 2018, Wen et al., 2024a) indicating that seizure phenotypes reflect systems-level transcriptional reprogramming.

Whole-brain RNA sequencing revealed a strongly directional transcriptional response dominated by downregulation of genes governing inhibitory neurotransmission, ion transport, axon guidance, cell adhesion, endothelial transport, and neuroactive ligand–receptor signalling. This pattern closely mirrors transcriptomic signatures reported in human epilepsy, including reduced expression of GABA receptor subunits, glutamate transporters, and potassium channels (Guelfi et al., 2019, Kjær et al., 2019). In *ap3b2* CRISPants, suppression of *gabra1/2, gabrb1/2/3, gabrg2, slc1a2/6*, and multiple *kcn* family members provides a molecular correlate for the hyperexcitable and hypersynchronous Ca²⁺ dynamics observed in vivo. Beyond neurotransmission, extensive downregulation of axon-guidance and neurodevelopmental pathways indicates disruption of early circuit assembly and refinement. Many DEE-associated genes converge on pathways regulating neuronal migration, axon growth, and synaptogenesis (Medyanik et al., 2025), and our findings suggest that AP3B2 loss perturbs developmental wiring processes, either resulting from, or independent of, synaptic vesicle trafficking. This developmental instability likely contributes to the persistent network-level hyperexcitability detected by Ca^²⁺^ imaging.

Importantly, since DEE is a genetically diverse but clinically identifiable umbrella disorder, we found many down regulated genes in *ap3b2* CRISPant brains that are associated with DEE. In total, 17 DEE-associated genes were identified by Enricher-GO analysis of the down regulated DEG list: *arhgef9* (DEE8), *cdk19* (DEE87), *gabra1* (DEE19), *gabra2* (DEE78), *gabrb1* (DEE45), *gabrb2* (DEE92), *gabrb3* (DEE43), *gabrg2* (DEE74), *grin2b* (DEE27), *hcn1* (DEE24), *kcna2* (DEE32), *kcnb1* (DEE26), *kcnh5* (DEE112), *kcnt1* (DEE14), *slc1a2* (DEE41), *slc25a22* (DEE3) and *synj1* (DEE53). This illustrates the underlying common causes of DEE and may mean that therapies found to work in one DEE may well work in others, even when no known direct link has been found.

### 4.4 Losartan provides partial rescue in two DEE models, suggesting potential for new approaches to treatment

Acute losartan treatment produced a reproducible but incomplete improvement in the *ap3b2* CRISPant phenotype. Behaviourally, losartan significantly reduced hyperlocomotion, and this was empowered by being able to use the same tadpoles before and after treatment. Due to the technical limitations of immobilising tadpoles long term, effects on individual Ca²⁺ event amplitude, frequency, and spectral power could only be compared between equivalent cohorts. While we observed a trend towards less overt Ca^2+^ events, the differences between groups and were not significant.

The relevance of losartan in this context lies in its capacity to modulate neurovascular and homeostatic pathways rather than directly targeting synaptic excitability. Losartan enhances thrombospondin-1–dependent activation of latent TGF-β (Bar-Klein et al., 2014), a signalling axis repeatedly implicated in seizure susceptibility in the setting of BBB dysfunction (Swissa et al., 2019). In the present study, transcriptomic suppression of multiple components associated with TGF-β responsiveness, including *tgfbr2* and *tgfbr3*, coincided with rapid sodium fluorescein leakage, indicating impaired BBB integrity. Rather than demonstrating overt neuroinflammation, our data support a state in which developing neural circuits are rendered more vulnerable to dysregulated vascular and immune influences. Losartan’s partial efficacy therefore could be explained by stabilisation of this permissive pathological environment, rather than correction of the primary synaptic trafficking defect caused by AP3B2 loss.

We previously showed that losartan was effective in reducing seizures in the *X. laevis neurod2* (DEE72) CRISPant tadpole model (Banerjee et al., 2024), suggesting that TGF-β–associated pathways represent a shared downstream vulnerability across genetically distinct DEEs. Our findings are also concordant with evidence from other epilepsy models and from clinical studies. In rodent seizure models, losartan reduces BBB permeability and seizure burden (Hong et al., 2019, Tchekalarova et al., 2016), while epidemiological studies report a reduced incidence of epilepsy among patients treated with angiotensin II receptor blockers, particularly losartan (Doege et al., 2022). At the same time, the absence of anticonvulsant effects in ex vivo human cortical tissue (Reyes-Garcia et al., 2019) underscores that losartan does not act as a conventional anti-seizure medication, and that its efficacy depends on engagement of intact neurovascular and immune-modulatory mechanisms.

Taken together, our data indicate that AP3B2-deficient networks retain sensitivity to pharmacological modulation of downstream regulatory pathways. Although the rescue observed here is partial and acute, the responsiveness to losartan identifies TGF-β–linked neurovascular mechanisms as functionally relevant modifiers of network instability in this DEE48 model. In this light, epidemiological evidence linking angiotensin II receptor blocker use, particularly losartan, to a reduced incidence of new-onset epilepsy (Wen et al., 2024b) provides independent support for the therapeutic relevance of targeting neurovascular and homeostatic pathways in epileptic encephalopathies.

## 5 Conclusions and Limitations of the study

This work demonstrates that loss of 80 to 85% of Ap3b2 in *X. laevis* tadpoles is sufficient to generate a robust DEE48-like phenotype, characterized by seizure-like behaviour, increased neural activity and interhemispheric synchrony and early blood–brain barrier leakage. Whole-brain transcriptomics revealed coordinated downregulation of inhibitory synaptic components, ion channels, axon-guidance pathways, and neuroinflammatory genes, providing a mechanistic framework that links AP3B2 deficiency to circuit instability and BBB fragility. The behavioural normalisation achieved with acute losartan treatment highlights the potential therapeutic relevance of targeting TGF-β–associated neuroinflammatory mechanisms. While our results support the use of *Xenopus* CRISPants as a rapid, integrative model for dissecting genetic DEE, several limitations should be noted. F₀ CRISPant tadpoles are both mosaic and carry varying levels of overall editing, with the variability on genotype likely contributing to variability in phenotype severity. Generating stable mutant lines would reduce such variability, but the severity of the human DEE phenotypes suggests raising and maintaining these could be challenging. RNA-seq was performed on whole brains, limiting cell-type resolution, and pooling of brains to create large enough samples could also reduce power. Losartan was assessed only acutely and at a single developmental stage and dose (albeit based on previous testing in another model), so the practicality of using it therapeutically in DEE remains untested. Finally, while *Xenopus* provides rapid access to early neurodevelopmental mechanisms, complementary studies will be essential to confirm the translational relevance of these findings.

## 6. Data Availability Statement

All data supporting the conclusions of this study are available from public repositories. Summary datasets and supporting analyses are provided in **Supplementary File 1.xlsx** and **Supplementary_Material.docx**. Raw and processed RNA-sequencing data have been deposited in the NCBI Gene Expression Omnibus (GEO) under accession **GSE312492**. The complete analysis pipeline is available via GitHub (https://github.com/sulagna-banerjee/xenopus-calcium-imaging-pipeline) and permanently archived on Zenodo (DOI: https://doi.org/10.5281/zenodo.17931981).

## 8. Author Contributions

Conceptualization: S.B., C.W.B., P.S.; Data curation: S.B., C.W.B.; Formal analysis and visualization: S.B., C.W.B., C.W.E, S.C.R.; Investigation S.B., C.W.E., C.W.B.; Funding acquisition, Project administration, Supervision: C.W.B., P.S.; Methodology: S.B., C.W.B., P.S. S.C.R; Writing-original draft preparation: S.B.,C.W.B, P.S. Writing-reviewing and editing: S.B., C.W.B., P.S.

## Acknowledgments

The authors thank Nikita Woodhead for *Xenopus* care, Joanna Ward for general lab technical assistance, Jack O’Neill for assistance designing the scrambled control sgRNA, and Edward Ruthazer and Anne Schohl for the kind gift of GCaMP6s-CS2 and mCherry-CS2+ plasmids.

