## Supplementary figure for "Seizures, increased interhemispheric synchrony, altered brain transcriptomics and a leaky blood-brain barrier result from loss of *ap3b2* in a CRISPR tadpole model of DEE48"

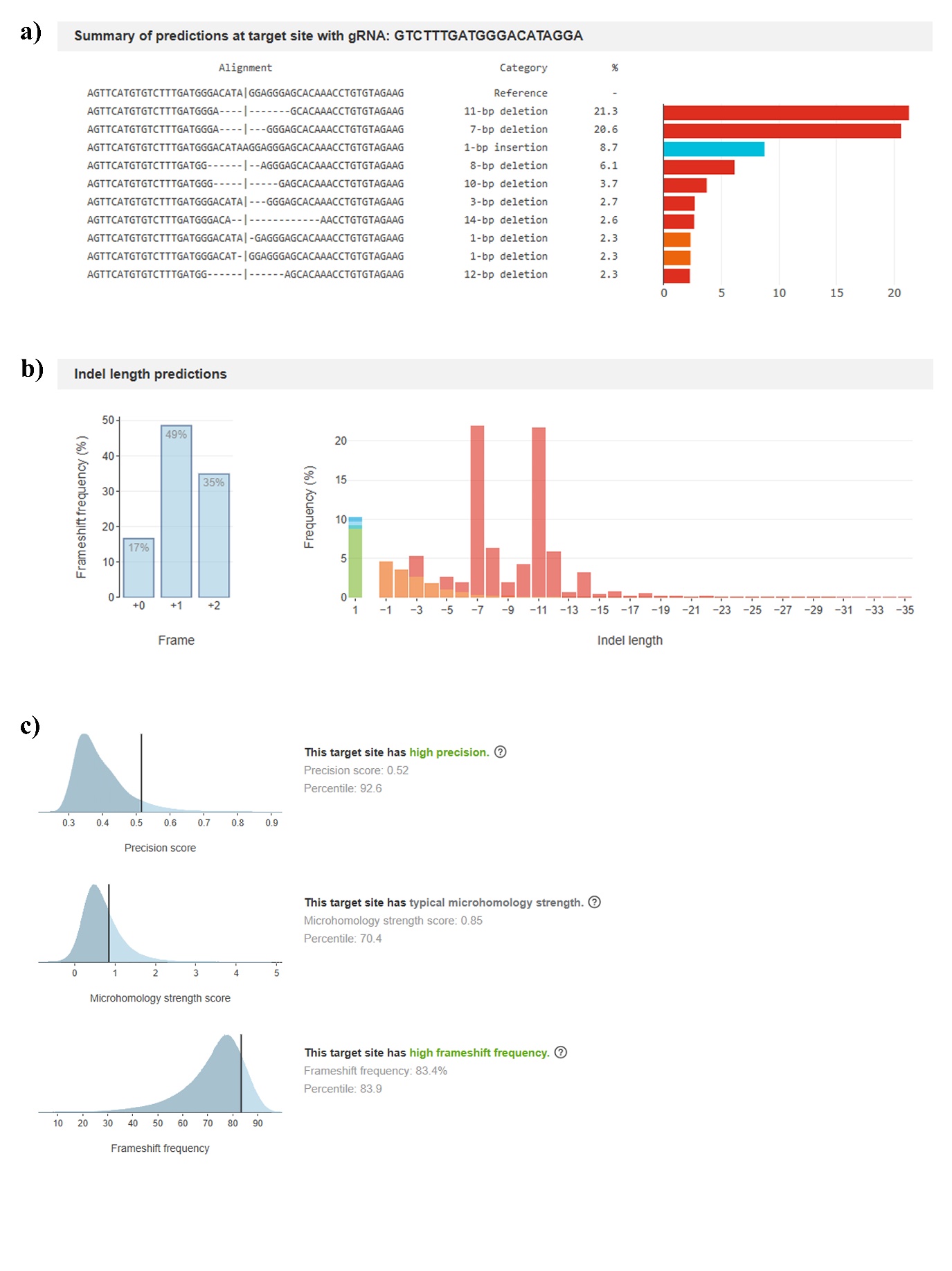

Figure S 1. InDelphi predictions ap3b2.S sgRNA 2.

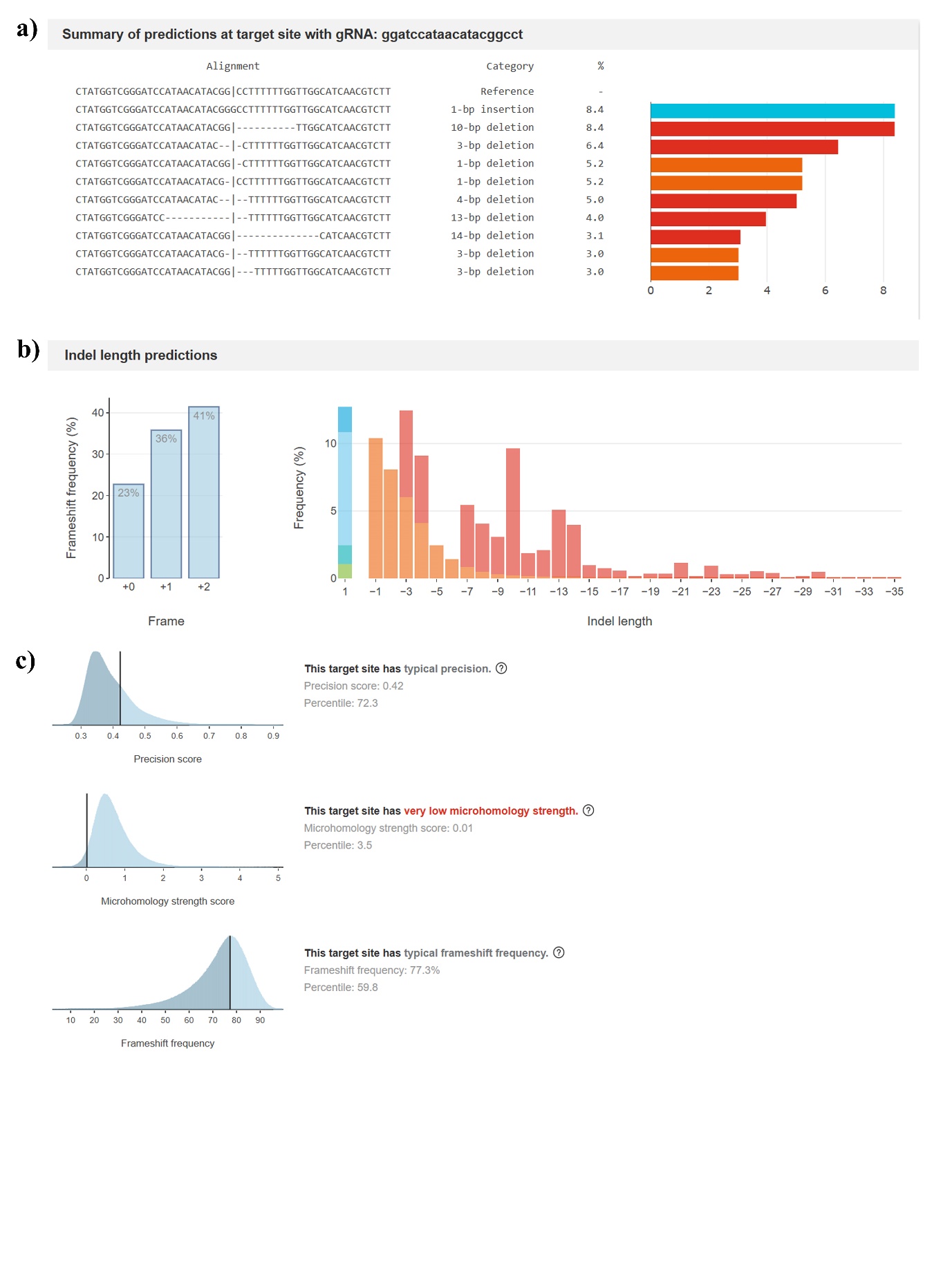

Figure S 2. InDelphi predictions ap3b2.S sgRNA 3.

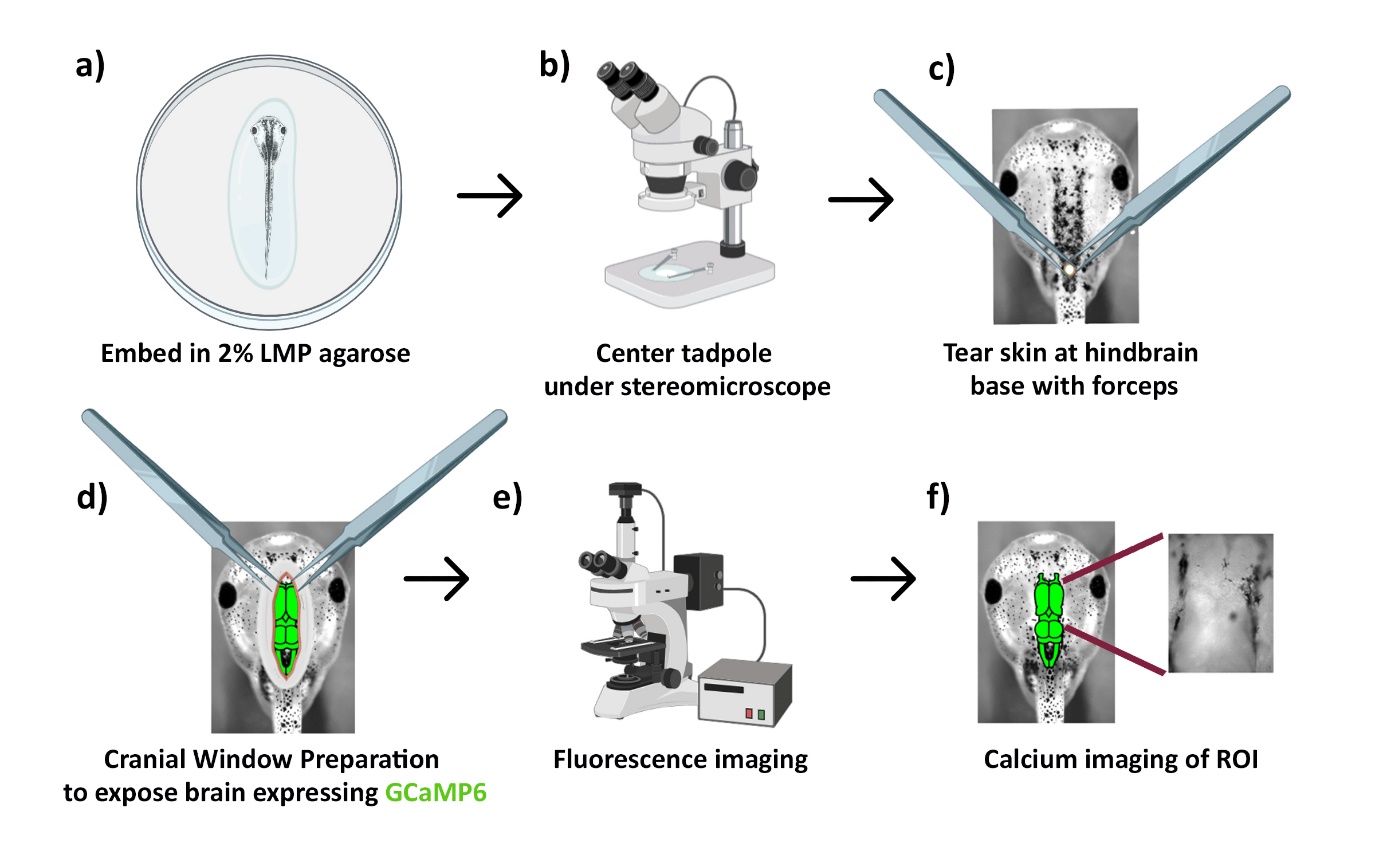

Figure S 3. Workflow for cranial window preparation and calcium imaging in Xenopus tadpoles. a) Tadpole embedded dorsal-up in 2% LMP agarose with nares/mouth free. b) Specimen centered under a stereomicroscope. c) Initial skin tear at the hindbrain base using fine forceps (Dumont #5). d) Cranial window preparation: skin is peeled anteriorly to expose the dorsal brain (hindbrain → midbrain → forebrain) expressing GCaMP6s. e) Transfer to the fluorescence microscope for imaging. f) Calcium imaging of defined brain regions (ROIs); inset illustrates a representative field used for time-series extraction. Arrows indicate the order of operations. Created in BioRender. Banerjee, S. (2026) <https://BioRender.com/1m35lxo>.

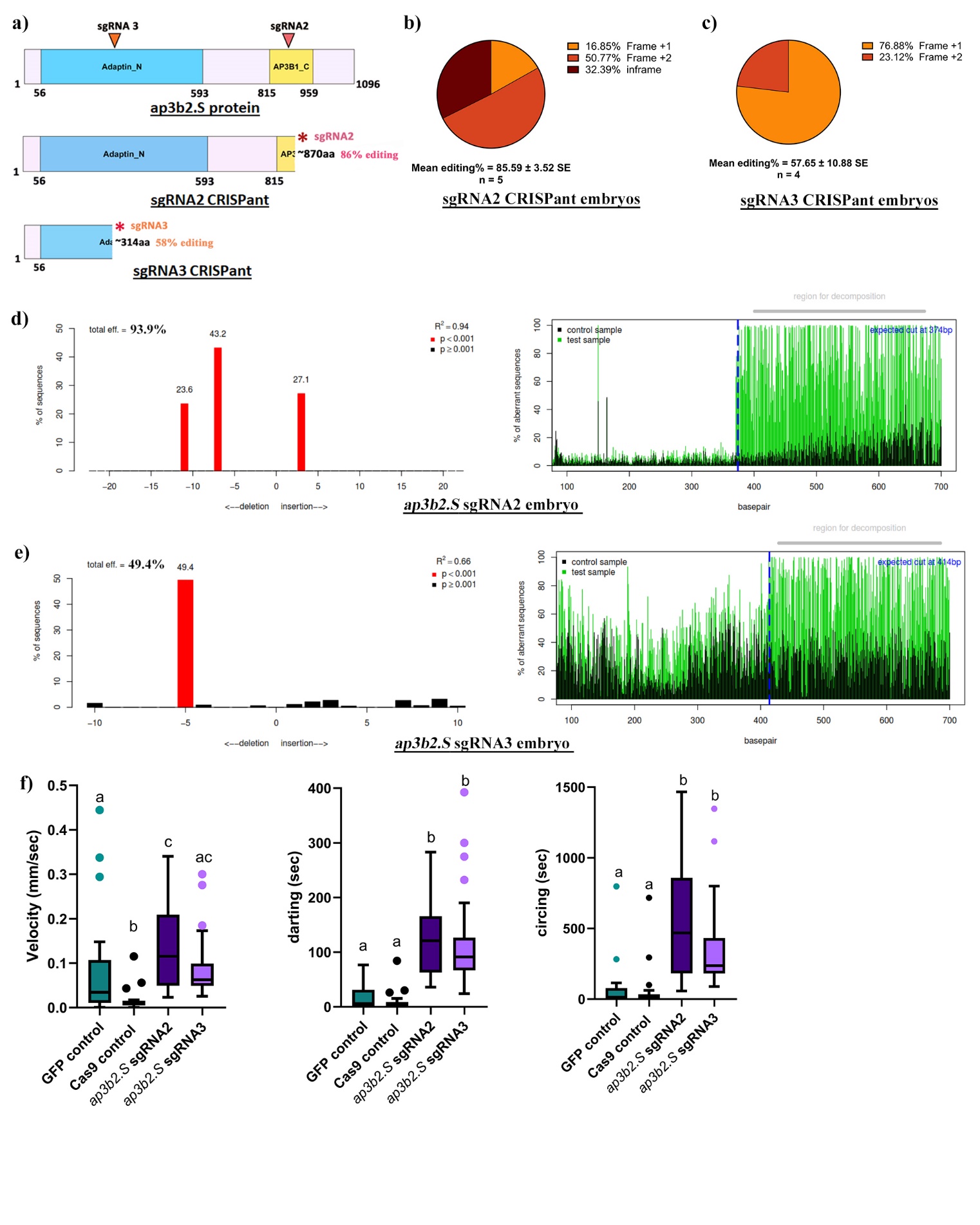

Figure S 4. Preliminary data showing predicted editing outcomes for both sgRNA2 and sgRNA3. (a) schematic of editing consequences. (b, c) editing summary. (d,e) example editing from TIDE (f) Behavioural data for two sgRNAs. N=24 for all groups. One way ANOVA with ad-hoc Kruskal-Wallis test. Raw data and statistics are in Supplementary File1.

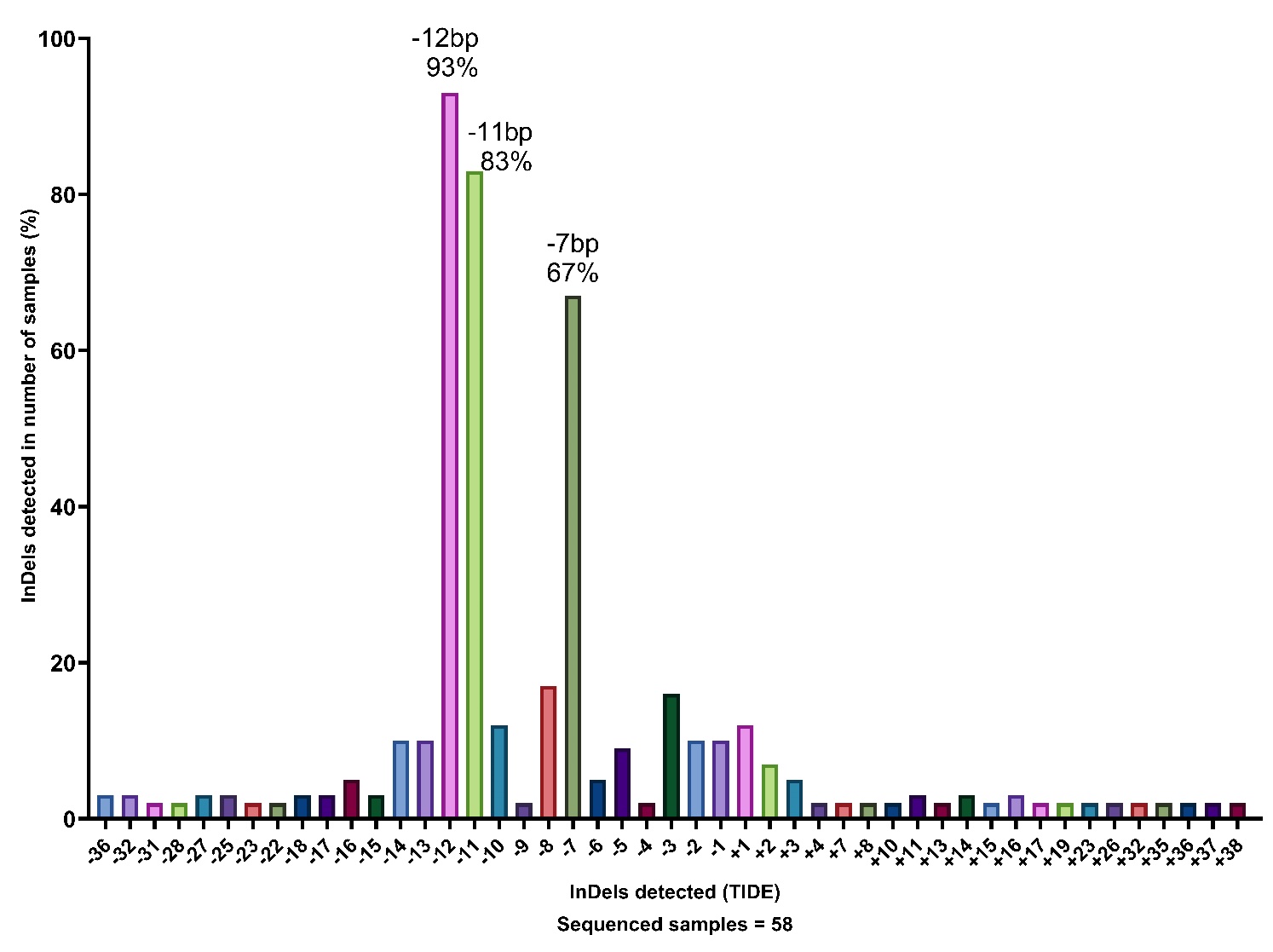

Figure S 5. Most detected InDels from all ap3b2.S sgRNA2 samples and TIDE analysis

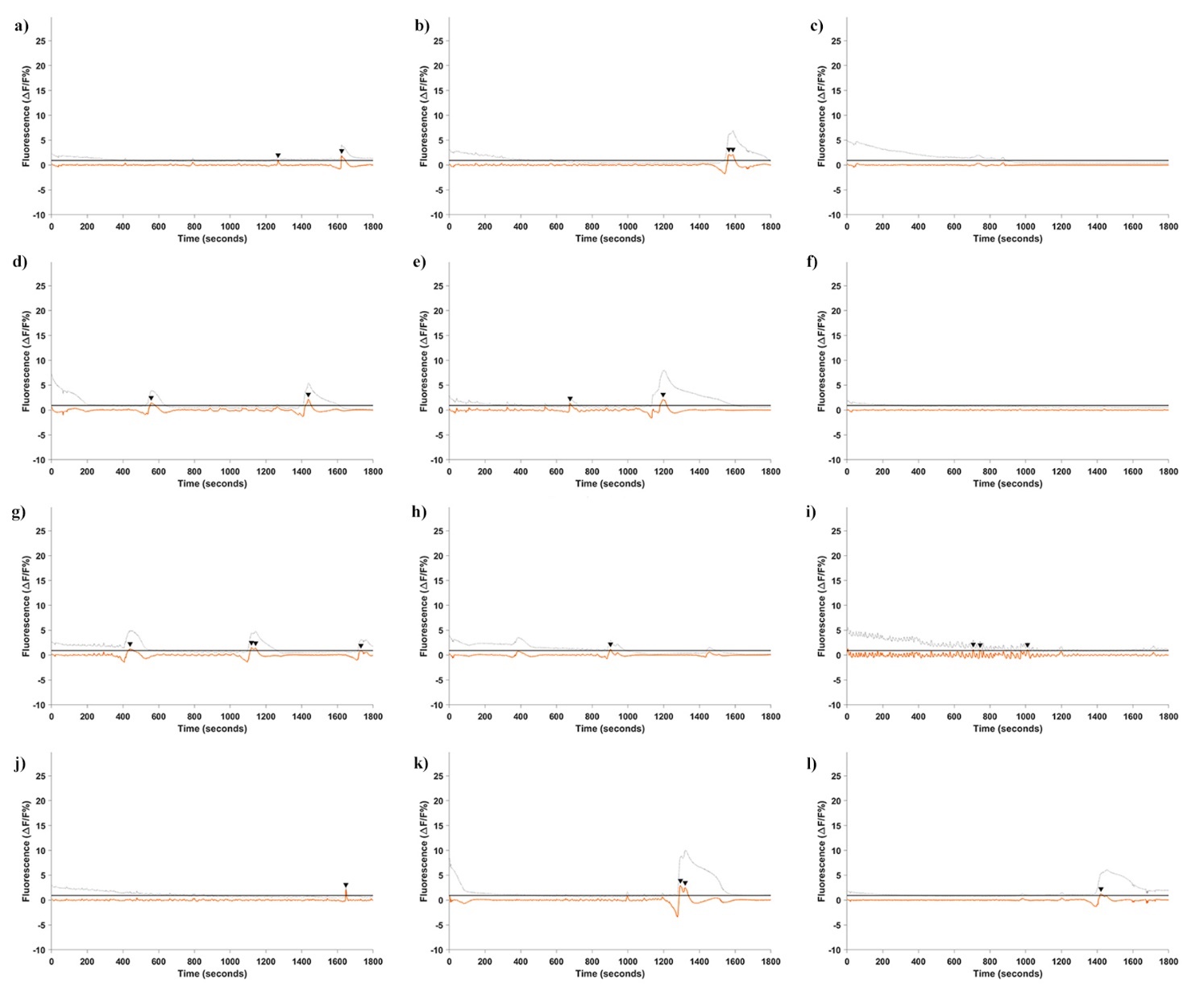

Figure S 6. Control tadpoles with no editing: all 12 Ca^2+^ baseline traces

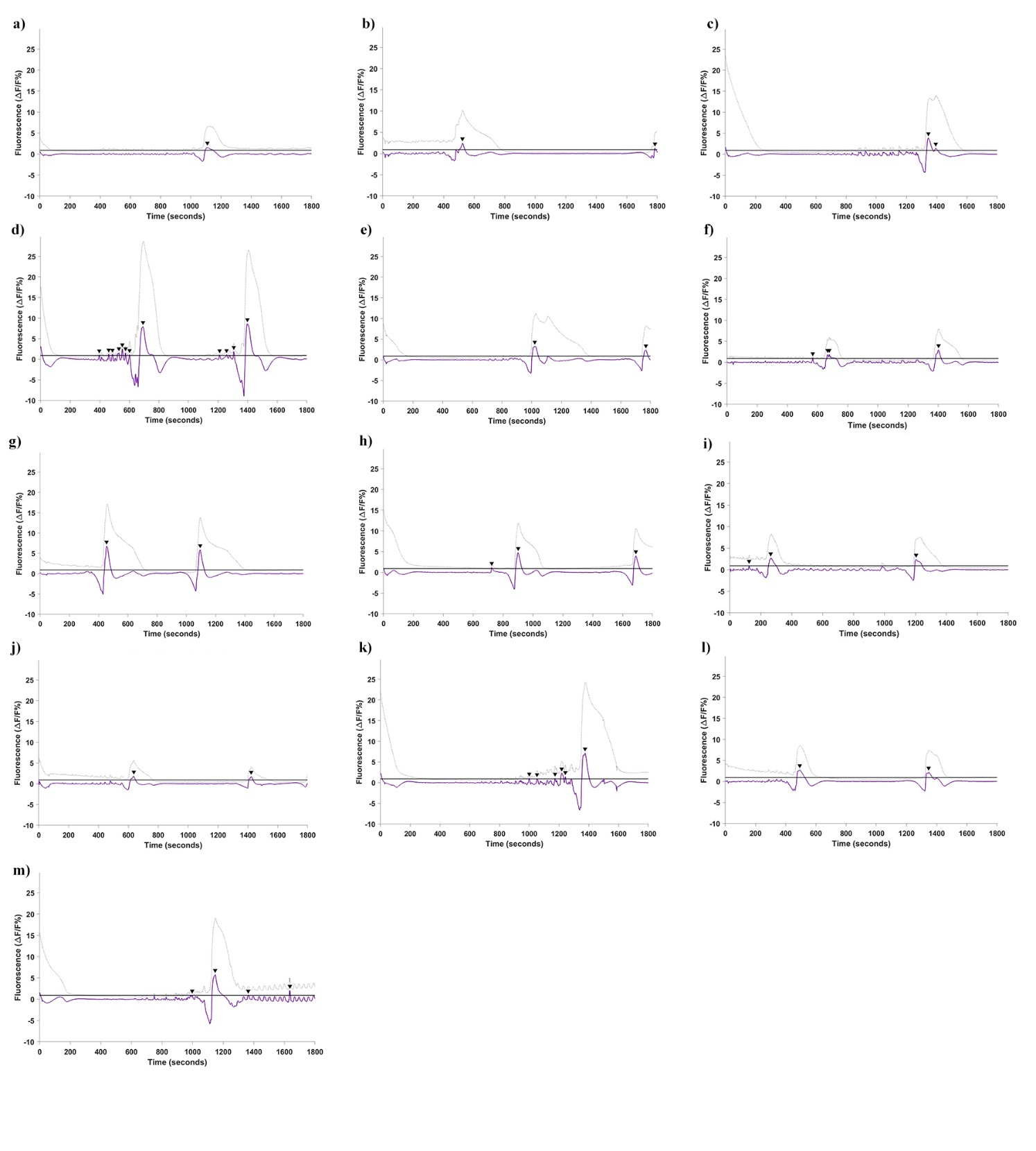

Figure S 7. All 13 tadpoles, *ap3b2* CRISPant baseline Ca^2+^ traces

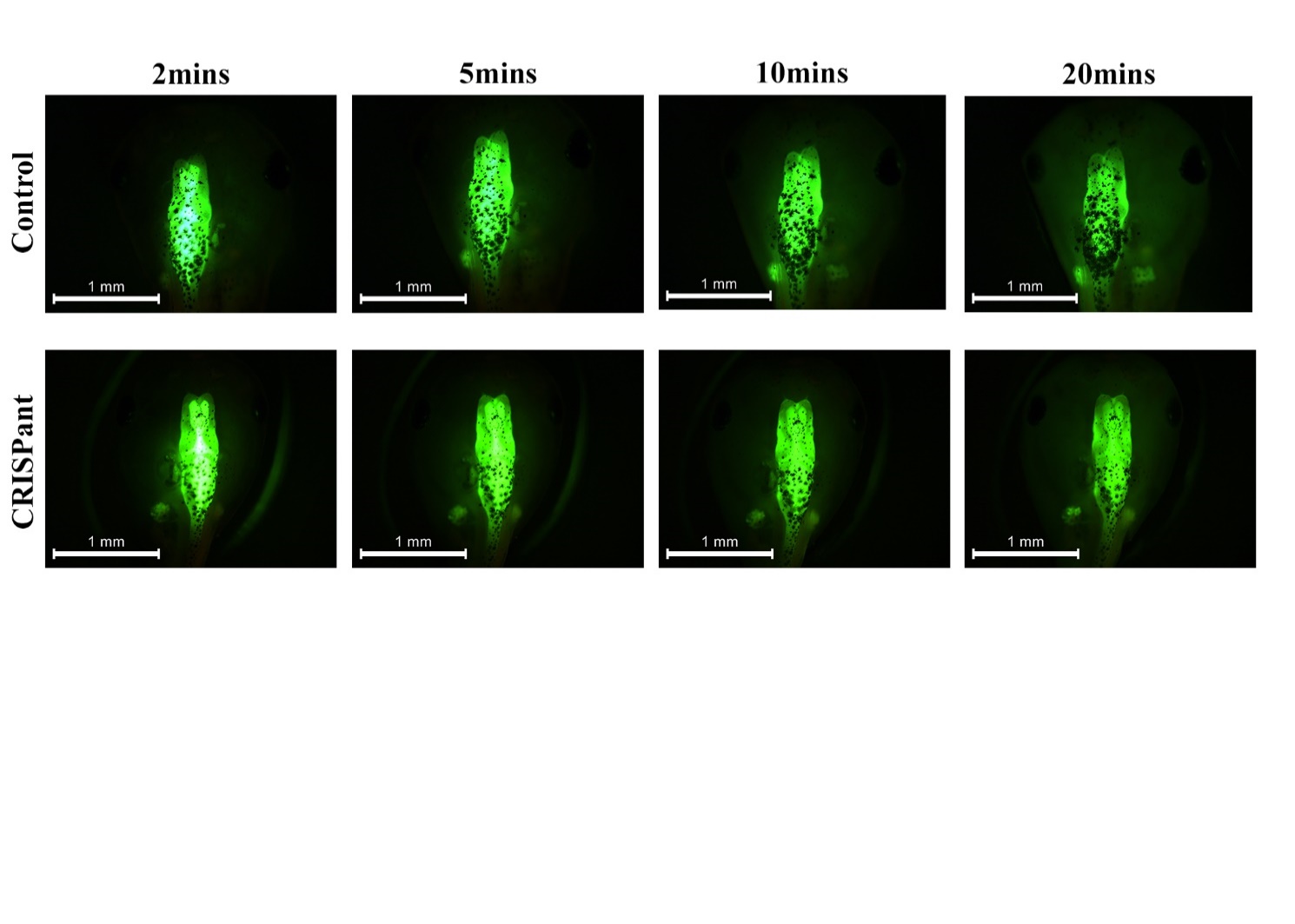

Figure S 8. BBB figs for all timepoints, example of typical control unedited and ap3b2 CRISPant images.

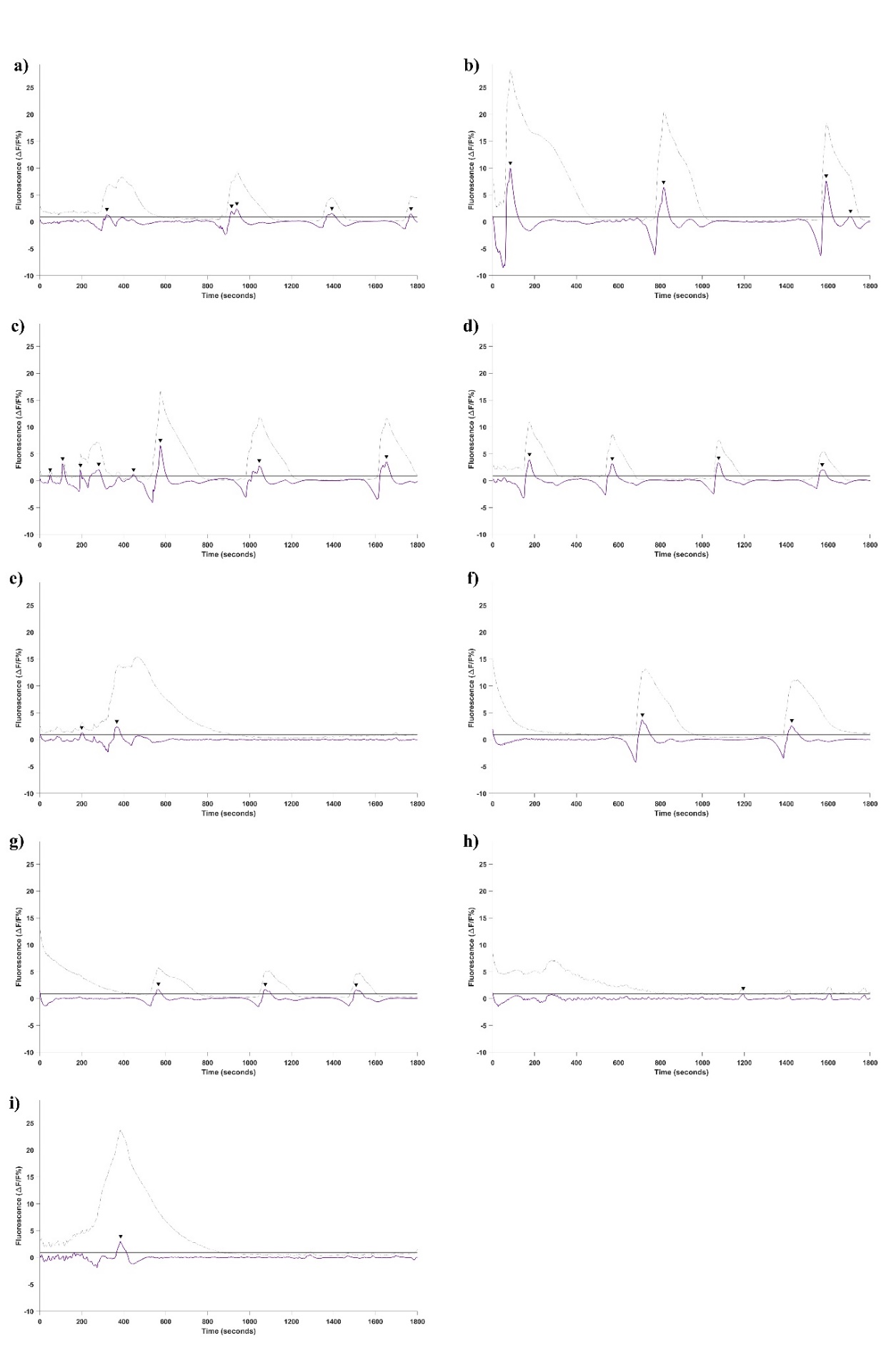

Figure S 9. All 9 untreated tadpoles, *ap3b2* CRISPant baseline Ca^2+^ traces (Losartan test)

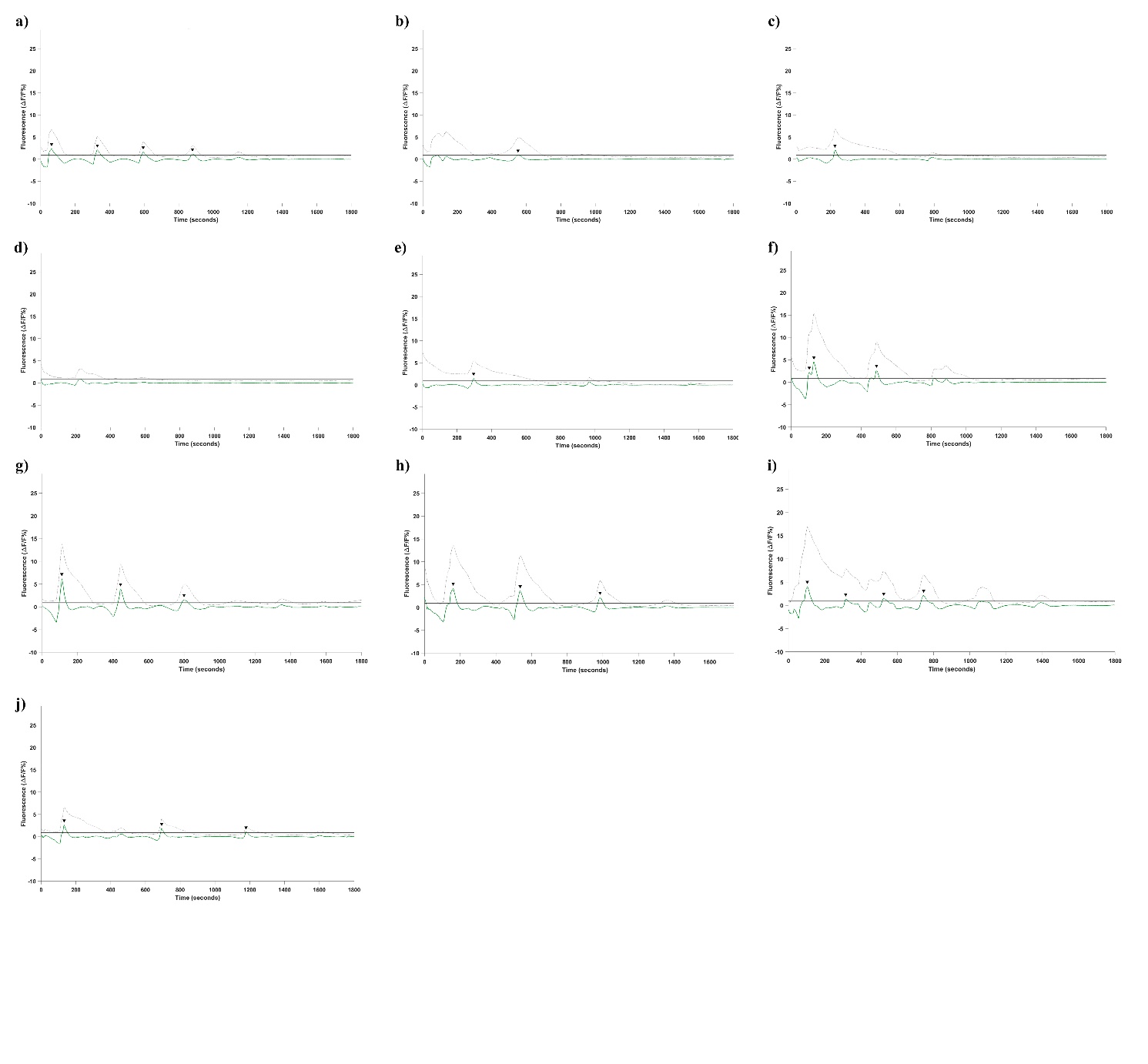

Figure S 10. All 10 losartan treated tadpoles, *ap3b2* CRISPant baseline Ca^2+^ traces (Losartan test)

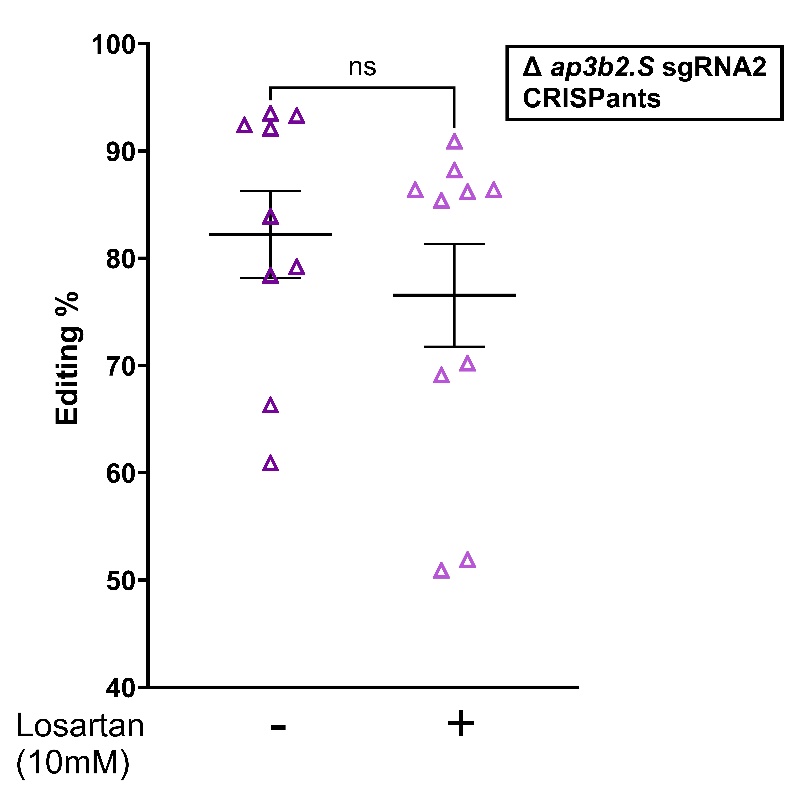

Figure S 11. Editing % comparison for Losartan calcium groups of tadpoles. Raw data and statistics in Supplementary File 1
